# BayesianSSA: a Bayesian statistical model based on structural sensitivity analysis for predicting responses to enzyme perturbations in metabolic networks

**DOI:** 10.1101/2024.03.14.585120

**Authors:** Shion Hosoda, Hisashi Iwata, Takuya Miura, Maiko Tanabe, Takashi Okada, Atsushi Mochizuki, Miwa Sato

## Abstract

**Background:** Chemical bioproduction has attracted attention as a key technology in a decarbonized society. In computational design for chemical bioproduction, it is necessary to predict changes in metabolic fluxes when up-/down-regulating enzymatic reactions, that is, responses of the system to enzyme perturbations. Structural sensitivity analysis (SSA) was previously developed as a method to predict qualitative responses to enzyme perturbations on the basis of the structural information of the reaction network. However, the network structural information can sometimes be insufficient to predict qualitative responses unambiguously, which is a practical issue in bioproduction applications. To address this, in this study, we propose BayesianSSA, a Bayesian statistical model based on SSA. BayesianSSA extracts environmental information from perturbation datasets collected in environments of interest and integrates it into SSA predictions.

**Results:** We applied BayesianSSA to synthetic and real datasets of the central metabolic pathway of *Escherichia coli*. Our result demonstrates that BayesianSSA can successfully integrate environmental information extracted from perturbation data into SSA predictions. In addition, the posterior distribution estimated by BayesianSSA can be associated with the known pathway reported to enhance succinate export flux in previous studies.

**Conclusions:** We believe that BayesianSSA will accelerate the chemical bioproduction process and contribute to advancements in the field.

## 1 Background

Chemical production using microbes, known as chemical bioproduction, has attracted attention as a key technology in a decarbonized society. Chemical bioproduction is expected to be essential for sustainable development [1], such as the production of medicines [2, 3], fuels [4], and foods [5], and the absorption of CO2 [6, 7]. For efficient chemical bioproduction, computational designs of metabolic networks are typically employed to reduce the cost of comprehensive wet lab experiments [8, 9].

One powerful strategy for computational design is to predict changes in metabolic fluxes when up-/down-regulating enzymatic reactions, that is, responses to enzyme perturbations [10, 11]. Up/down-regulation of enzymatic reactions through genetic manipulations, such as modification, overexpression, and knockout [12, 13], can alter the metabolic fluxes. Increasing the flux of the target chemical leads to efficient chemical bioproduction, and prediction is an essential step in this process.

There are two types of computational methods for predicting responses to enzyme perturbations. The first is flux balance analysis (FBA)-based methods [14, 15, 16, 17], which maximize an objective function, instead of considering kinetics. When applied to unfamiliar strains, FBA-based methods suffer from dependence on the objective function, which is typically the biomass objective function [18]. Specifically, extensive wet lab experiments need to be conducted to measure phenotypes, such as biomass, of the strain of interest and to identify the objectives of the strain in biological activities [19]. The second is kinetics-based methods, such as sensitivity analysis in kinetic models [20, 21, 22] and structural sensitivity analysis (SSA) [10, 23]. Sensitivity analysis in kinetic models allows predicting quantitative responses to enzyme perturbations. There are global and local sensitivity analysis [24, 22], and the local sensitivity analysis is the more typical method for applying to kinetic models, which is called metabolic control analysis [25] and has been successfully used in many applications [26, 27, 28]. To construct a kinetic model, it is necessary to obtain parameters and functional forms of reaction rates of all reactions in the metabolic network of interest under specific environmental conditions [20, 21]. Therefore, parameter estimation is typically essential for unfamiliar strains, whose known information is rarely available [29]. Even though many parameter estimation methods have been developed [30, 31, 32, 33], it can still be a cumbersome process due to the high dimensionality of the parameter space [30, 34]. In contrast, SSA can predict qualitative responses, which are the signs of responses, to enzyme perturbations only from structural information of the metabolic network. SSA does not need to determine the functional forms and parameters of the reaction rates. In other words, SSA removes the burden of parameter determination entirely. Because of its parameter-free nature, SSA could be widely applicable to chemical bioproduction.

SSA is originally a method to predict qualitative responses of a chemical reaction system to perturbations in enzyme amounts (or activities) only from structural information of the reaction network. In SSA, change in the chemical concentrations/fluxes to a perturbation is given in the form of a rational function of “SSA variables,” which are defined as derivatives of reaction rates with respect to chemical concentrations. From their definitions (*cf*. the “Our model” section), the SSA variables vary depending on environmental conditions, such as aerobic/anaerobic states, pH levels, and nutrient availability, as well as individual differences among microbes. By the SSA theory, whether the rational function is zero or non-zero, indicating the absence or presence of a response, is unambiguously determined from the structural information of the network alone. Furthermore, the sign of the rational function, indicating a positive or negative response, could potentially be determined by considering the signs of SSA variables, which are usually deducible from general considerations or biological knowledge. However, if the range of the rational function happens to include zero, its sign becomes undeterminable. For example, when a rational function is a subtraction of two positive variables, its sign depends on the quantitative values of these two variables. We refer to such structurally undeterminable response predictions as “indefinite predictions.” Indefinite predictions are true as the general consequences of responses drawn from structural information alone, which SSA aims to obtain rather than specific consequences.

However, these indefinite predictions in SSA can present practical challenges in the application to specific species and culture environments for chemical bioproduction, due to the necessity of precisely detecting reactions to up-/down-regulate. They often arise in complex and intertwined metabolic networks. Indeed, the central metabolic pathway analyzed in this study exhibits many indefinite predictions (Supplementary Table S1). As mentioned above, the SSA variables vary depending on the environmental conditions, fluctuating in accordance with the individual differences among microbes. For specific microbial species and culture environments, constraining the possible values of SSA variables on the basis of environmental information may decrease the number of indefinite predictions. This implies that application methods of SSA using environmental information are practically beneficial for chemical bioproduction.

In this study, we propose BayesianSSA, a Bayesian statistical model that extracts environmental information from perturbation datasets collected in environments of interest and integrates it into SSA predictions. BayesianSSA considers the variables of SSA as stochastic variables, and they are estimated using the perturbation data. Although BayesianSSA introduces new parameters to estimate, it still requires fewer parameters per reaction than kinetic modeling requires. For example, for a one-substrate reaction, BayesianSSA requires one parameter while kinetic modeling with the Michaelis–Menten equation requires two parameters, *V*_max_ and *K*_m_. In addition, BayesianSSA does not need to explore the functional forms of the reaction rates. The introduction of stochastic variables in BayesianSSA brings two additional advantages. First, it allows for the consideration of the uncertainty caused by individual differences among microbes. The variables of SSA may depend not only on environmental conditions but also on individual differences among microbes, and considering their uncertainty may contribute to predictive performance. Second, it enables a probabilistic interpretation of indefinite prediction by positivity confidence values. These values are defined as the probability that the predictive response is positive. We report the results of applying BayesianSSA to synthetic and real datasets of the central metabolic pathway of *E. coli*. To validate the practicability of BayesianSSA and assess whether BayesianSSA can integrate environmental information into SSA predictions, we compared predictive performances between BayesianSSA and a base method, which is the same as the BayesianSSA model but utilizes an initial prior distribution without incorporating perturbation datasets, as well as a naive Bayes model on these datasets. Utilizing environmental information may enhance predictions for out-of-sample perturbations, where chemical reactions to be perturbed are not included in the sample used to fit BayesianSSA for a given target chemical. To examine this effect, we evaluated predictions of out-of-sample perturbations by BayesianSSA.

## 2 Theoretical background

### 2.1 Structural sensitivity analysis

#### 2.1.1 Algorithm

We explain the SSA algorithm [10, 11] briefly in this section. Consider the following ordinary differential equation (ODE):

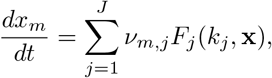

where *x*_*m*_ is the concentration of the *m*-th metabolite, *J* is the number of different reactions, *ν*_*m,j*_ is the (*m, j*) element of the stoichiometric matrix ***ν*** that indicates metabolites as rows and reactions as columns, *F*_*j*_(*·*) is the *j*-th reaction rate function, which shows the reaction flux, *k*_*j*_ is the reaction rate constant of the *j*-th reaction, **x** = (*x*_1_, …, *x*_*M*_)^T^ is the vector of metabolite concentrations, and *M* is the number of different metabolites. Under the situation where the system obeys this ODE and the assumption that *F*_*j*_ monotonically increases with respect to *k*_*j*_, the SSA algorithm can derive the rational function of a response to a perturbation of the *j*-th reaction. Here, the perturbation and response are represented as an operation changing *k*_*j*_ to *k*_*j*_ + *δk*_*j*_ [11] and the change in metabolite concentrations/reaction fluxes from the initial steady-state to the eventual steady-state after the perturbation, respectively.

The first step of the SSA algorithm is to make a matrix **R**(**r**). The (*j, m*) element of **R**(**r**), denoted by *r*_*j,m*_, is equal to *∂F*_*j*_*/∂x*_*m*_. We write non-zero elements of **R**(**r**) collectively as **r** ∈ ℝ^*P*^, with *P* being the total number of non-zero values in **R**(**r**). That is, the matrix **R**(**r**) indicates the dependence of reactions to metabolites. For example, *r*_*j,m*_ *>* 0 if the *m*-th metabolite is a substrate of the *j*-th reaction and *r*_*j,m*_ = 0 otherwise. Using the matrix **R**(**r**), the matrix **A**(**r**) ∈ ℝ^(*J*+*L*)*×*(*M*+*K*)^ is defined

As

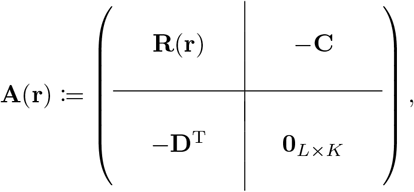

where **0**_*L×K*_ ∈ ℝ^*L×K*^ is a zero matrix, whose elements are all zero, **C** = (**c**_1_, …, **c**_*K*_) ∈ ℝ^*J×K*^, **c**_*k*_ is the *k*-th basis of ker ***ν***, which indicates the right null space of ***ν***, *K* is the number of the bases of ker ***ν*, D** = (**d**_1_, …, **d**_*L*_) ∈ ℝ^*M×L*^, **d**_*l*_ is the *l*-th basis of ker ***ν***^T^, which indicates the right null space of ***ν***, and *L* is the number of the bases of ker ***ν***^T^. Note that **A**(**r**) is in general a square matrix, *i*.*e*., *J* + *L* = *M* + *K*. A sensitivity matrix is defined as

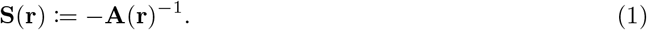

Here, the (*m, j*) element of **S**(**r**) ∈ ℝ^(*M*+*K*)*×*(*J*+*L*)^ is proven to be equal to a constant multiple of a quantitative response of the *m*-th metabolite concentration/flux to a perturbation of the *j*-th reaction/conserved quantity [11]. Even though we calculate the sensitivity matrix, we cannot determine the quantitative response value. However, each element of the sensitivity matrix has the same sign (positive, negative, or zero) as the corresponding quantitative response value, enabling us to discuss qualitative responses. Although flux responses are represented as sets of reactions that are non-zero in **c**_*k*_, we can obtain a flux response corresponding to each reaction by calculating linear combinations. Let **T**(**r**) ∈ ℝ^*J×*(*J*+*L*)^ be a matrix indicating the responses of each reaction flux, which can be written as

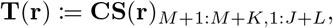

where **S**(**r**)_*M*+1:*M*+*K*,1:*J*+*L*_ ∈ ℝ^*K×*(*J*+*L*)^ is a block matrix of **S**(**r**), which is extracted from the (*M* + 1)th to (*M* + *K*)-th rows.

#### 2.1.2 Qualitative response prediction

SSA can predict qualitative responses (positive, negative, zero, or indefinite) to perturbations using only structural information, which is the metabolic network and the constraints on the **r** values. Here, we describe this in a precise way.

Let **Q**(**r**) ∈ {−1, 0, 1}^(*M*+*J*)*×*(*J*+*L*)^ be the qualitative response matrix, which is given by

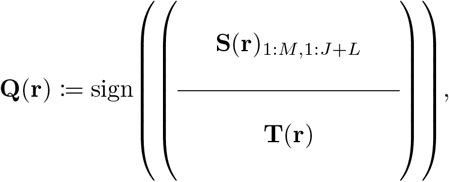

where sign(*·*) is the element-wise sign function, which returns a matrix whose element equals 1, 0, and –1 if the sign of the corresponding element of the given matrix is positive, zero, and negative, respectively. The responses of the *m*-th metabolite concentration and the *j*-th reaction flux to perturbations correspond to the *m*-th and (*M* + *j*)-th rows of **Q**(**r**), respectively. We refer to the *m*-th row and the *j*-th column of **Q**(**r**) as the *m*-th “observation target” and the *j*-th “perturbation target”, respectively. In addition, we call the perturbation experiment/prediction/response for the *m*-th observation and *j*-th perturbation targets the (*m, j*) experiment/prediction/response.

Since SSA is concerned with general results of qualitative responses, the elements of **Q**(**r**) are examined across all possible values of **r**. It is important to note that there are typically constraints on the **r** values, such as *r*_*j,m*_ *>* 0, and **Q**(**r**) is evaluated within these constraints. Let *q*_*m,j*_(**r**) denote the (*m, j*) element of **Q**(**r**) (*m* = 1, …, (*M* + *J*), *j* = 1, …, (*J* + *L*)). The qualitative response for each *q*_*m,j*_(**r**) can be classified into one of four categories; i) *q*_*m,j*_(**r**) is zero for any **r**. This case is a consequence of structural properties, as explained by a theorem known as the law of localization and buffering structures [35, 11]. ii) *q*_*m,j*_(**r**) is positive for any **r**. iii) *q*_*m,j*_(**r**) is negative for any **r**. iv) *q*_*m,j*_(**r**) varies depending on the quantitative values of **r**, making the sign of the response indefinite. For example, including a term *r*_1_ − *r*_2_ in the symbolic expression corresponding to *q*_*m,j*_(**r**) makes *q*_*m,j*_(**r**) positive if *r*_1_ *> r*_2_ and negative if *r*_1_ *< r*_2_, where *r*_*i*_ is the *i*-th element of **r**. These four categories are referred to as zero, positive, negative, and indefinite, respectively.

There are several methods for evaluating *q*_*m,j*_(**r**) for all possible **r**. One approach is to perform symbolic calculations, regarding **r** as symbolic variables [10]. Although this method is rigorous, it is only practical for relatively small metabolic networks due to its computational complexity. An alternative, more computationally tractable method is to draw a sufficient number of **r** samples from an arbitrary probabilistic distribution and numerically evaluating **Q**(**r**) [36]. This latter approach has inspired us to introduce stochastic variables into SSA in this study.

## 3 Methods

### 3.1 Our model

In SSA, the predictive response is obtained by considering all cases of **r. r** depends on {*k*_*j*_}_*j*_ and {*x*_*m*_}_*m*_, which, in turn, depend on environmental conditions, such as aerobic/anaerobic states, pH levels, and nutrient availability, as well as individual differences among microbes. The SSA qualitative response prediction described in the previous section thus focuses on the general consequences of responses drawn from structural information alone. We propose a Bayesian statistical model based on SSA, named BayesianSSA, to integrate environmental information extracted from perturbation data into SSA predictions. We consider a probability of each **r** value, regarding **r** as a stochastic variable. The BayesianSSA posterior probability of **r** reflects perturbation data and thus extracts environmental information. Predicting responses on the basis of the posterior distribution of **r** and positivity confidence values enables us to integrate the environmental information into SSA predictions. A positivity confidence value, which we proposed in this study, indicates the probability that the predictive response is positive. The positivity confidence values enable us to interpret indefinite predictions stochastically. We will describe the details in the “Positivity confidence value” section.

#### 3.1.1 Prior distribution and likelihood

In this section, we describe the likelihood function and the prior distribution of BayesianSSA. Suppose that we have a perturbation dataset **y**, which shows the signs of experimentally observed responses. We define the perturbation record, an element of **y**, as follows:

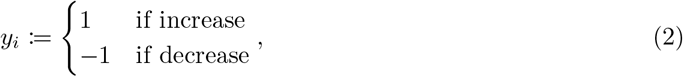

where *y*_*i*_ is the *i*-th perturbation record, obtained from the (*m*_*i*_, *j*_*i*_) experiment, and “increase” and “decrease” indicate the cases where the observation target increases and decreases when the perturbation target is perturbed, respectively (see the “Qualitative response prediction” section for the definition of observation and perturbation targets). We consider the probability that 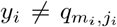(**r**) because experimental errors may occur even if *q*_*m,j*_(**r**) makes the correct prediction, assuming that the likelihood function is the following:

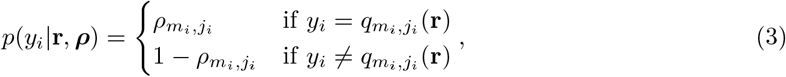

where *ρ*_*m,j*_ ∈ (0, 1) is a parameter indicating the reliability of the (*m, j*) experiment, and ***ρ*** is a matrix whose (*m, j*) element is *ρ*_*m,j*_. The likelihood function means that the probability of *y*_*i*_ is 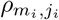 if **r** can accurately predict *y*_*i*_, and is 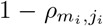 otherwise. In other words, BayesianSSA assumes that the result of the *i*-th experiment can stochastically vary in accordance with the probability of 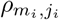 is different for each (*m, j*), and each (*m, j*) experiment is assumed to have different reliability in BayesianSSA. This assumption is reasonable because the distribution of measured values is supposed to be different for each (*m, j*) experiment.

We consider the prior distribution of *ρ*_*m,j*_ as

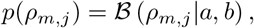

where *ℬ* (*·*|*a, b*) denotes the beta distribution probability density function with parameters *a* ∈ ℝ_*>*0_ and *b* ∈ ℝ_*>*0_. The prior distribution of **r** should be chosen in accordance with the constraint on **r**. A typical constraint on *r*_*j,m*_ is *r*_*j,m*_ *>* 0 with the *m*-th metabolite being the substrate of the *j*-th reaction, and we consider only such type of constraints in this study. We used a weighted empirical distribution with samples drawn from a log-normal distribution as the prior distribution. Specifically, t he prior distribution of **r** is the following:

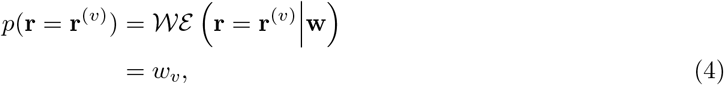

where *𝒲 ℰ* (**r** = **r**^(*v*)^∣**w)** denotes the weighted empirical distribution probability mass function with a stochastic variable **r**, a weight parameter **w** = (*w*_1_, …, *w*_*V*_)^*T*^ and a parameter sample set 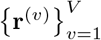 and 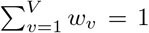, and *V* is the size of the sample set. Here, we generated the parameter sample set 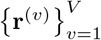 as follows:

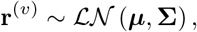

where *ℒ 𝒩* (***µ*, Σ**) denotes the log-normal distribution with parameters ***µ*** ∈ ℝ^*P*^ and **Σ** ∈ ℝ^*P ×P*^ (***µ*** and **Σ** correspond to the mean and covariance matrix parameter of the normal distribution, respectively), and *a* ∼ *𝒟* indicates that a stochastic variable *a* is drawn from a distribution *𝒟*.

The purpose of BayesianSSA is to obtain a better distribution *p*(**r**) for calculating positivity confidence values (Eq. (7)). Therefore, we need the marginal posterior distribution *p* (**r**|**y**). The marginal likelihood is obtained as

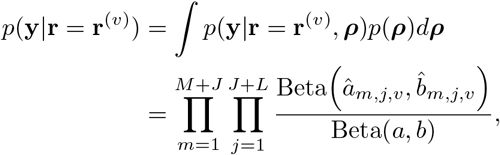

where Beta(*·, ·*) is the beta function, 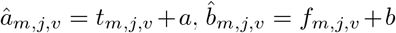, and *t*_*m,j,v*_ and *f*_*m,j,v*_ are the number of true and false (*m, j*) predictions based on **r**^(*v*)^ in **y**, respectively. Here, *t*_*m,j,v*_ = *f*_*m,j,v*_ = 0 for a (*m, j*) experiment that has not been conducted.

If a continuous prior and posterior distribution is desired, one can use them and obtain samples from the posterior distribution using the Markov chain Monte Carlo (MCMC) method. Since we discretized the log-normal distribution (Eq. (4)), we can calculate the posterior distribution without approximation using MCMC (details are described in the “Calculating posterior distribution” section).

#### 3.1.2 Calculating posterior distribution

The marginalized posterior distribution *p*(**r**|**y**) in BayesianSSA can be calculated by normalizing the following formula:

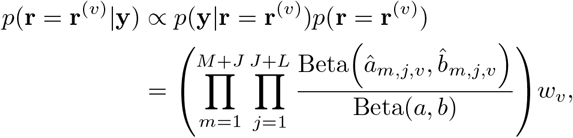

we can omit calculating the constant of this formula, and we derive that

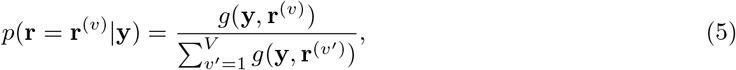

where

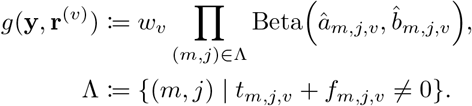

we implemented these calculations using the log-sum-exp trick.

#### 3.1.3 Calculating predictive distribution

To evaluate predictive performance, we calculate the predictive probability *p*(**y**^new^|**y**), where **y**^new^ is a new sample that is not used in BayesianSSA fitting. U sing t he p osterior d istribution *p*(**r, *ρ***|**y**) (Supplementary Section S1), the predictive probability can be obtained as

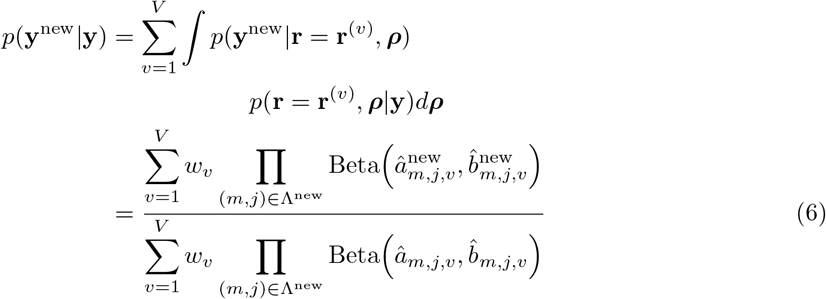

where

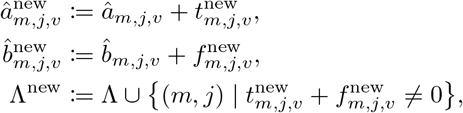

and 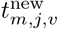 and 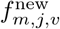 are the number of true and false (*m, j*) predictions based on **r**^(*v*)^ in **y**^new^, respectively. The detailed derivation is described in Supplementary Section S2.

#### 3.1.4 Bayesian updating

Bayesian updating, a procedure where the posterior distribution is used as a prior distribution for the next estimation, can be easily applied to BayesianSSA. In BayesianSSA, updating *â*_*m,j,v*_ and 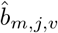 to 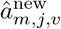 and 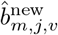 is equivalent to Bayesian updating (Supplementary Section S3). This update is also derived from the fact that the final updated posterior distribution in Bayesian updating does not depend on the order of the given perturbation data due to Bayes’ theorem [37].

#### 3.1.4 Positivity confidence value

The introduction of stochastic variables **r** enables interpreting indefinite predictions in SSA. We define positivity confidence values as the probabilities that the responses are positive for each indefinite prediction. The positivity confidence value of the (*m, j*) prediction for a distribution *p*^⋆^(**r**) is written as

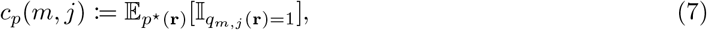

where 𝔼_*p*_^⋆^ (**r**)[*·*] denotes the expectation with respect to *p*^⋆^ (**r**), and 𝕀_*a*=*b*_ is an indicator that is equal to 1 if *a* = *b* and 0 otherwise. A higher positivity confidence value indicates that the qualitative response is more likely to be positive, and the probability of being negative is higher than that of being positive when the positivity confidence value is below 0.5. Note that we use the predictive distributions derived in the “Calculating predictive distribution” section rather than positivity confidence values to evaluate predictive performance on perturbation datasets. This is because positivity confidence values do not consider experimental error.

### 3.2 Used data

#### 3.2.1 Metabolic network information

We utilized the central metabolic pathway of *E. coli* MG1655 used in a previous study [38]. This metabolic network was originally from the EcoCyc database [39] and modified by Trinh *et al*. [40] and Toya *et al*. [38]. We preprocessed this dataset as follows:

1. Remove the biomass objective function.
2. Remove metabolites that have no reactions that produce or use them.
3. Remove reactions that no longer have substrates or products due to the previous processes.
4. Integrate cytoplasm and extracellular metabolites.

The second step is necessary to apply SSA to the network because it is typically impossible to calculate the inverse of the matrix **A**(**r**) (Eq. (1)) when metabolites not involved in the flow are included in the network. Here, metabolites in flow refer to metabolites that serve as both inputs and outputs of reactions included in the network. The fourth step aims to reduce computation time, and this procedure does not alter the results from SSA. After these preprocessing steps, we converted the resulting network into the stoichiometric matrix ***ν***. Constraints on **R**(**r**) were determined solely by the metabolic network, where we set *r*_*j,m*_ *>* 0 if the *j*-th metabolite is a substrate of the *m*-th reaction and *r*_*j,m*_ = 0 otherwise. The resulting metabolite and reaction lists are shown in Supplementary Table S2 and Supplementary Table S3. We use the abbreviations in Supplementary Table S2 and Supplementary Table S3 in the following.

#### 3.2.2 Synthetic data generation

We generated synthetic data to compare BayesianSSA with random and base methods. The synthetic perturbation dataset is generated in accordance with a Bernoulli distribution with parameters generated in accordance with beta distributions. Then, we replaced all 0 with −1 to match the support of the Bernoulli distribution and the range of the perturbation record *y*_*i*_ (Eq. (2)). We used GND, PTS, and PPC as the perturbation targets and succinate export (SUCCt) as the observation target. We used a beta distribution with an expectation is close to 0 for GND and PTS and a beta distribution with an expectation is close to 1 for PPC. These distribution settings were in accordance with previous reports [41, 42, 43, 44]. We used (*a, b*) = (0.1, 0.3) and (*a, b*) = (0.3, 0.1) as parameters of beta distributions with expectations are close to 0 and 1, respectively. Note that these parameters are used only for the data generation and different from the parameters of the prior distributions, which are weakly informative. We used 30 as the number of records for each perturbation target. The obtained synthetic dataset is shown in Table 1.

**Table 1:**
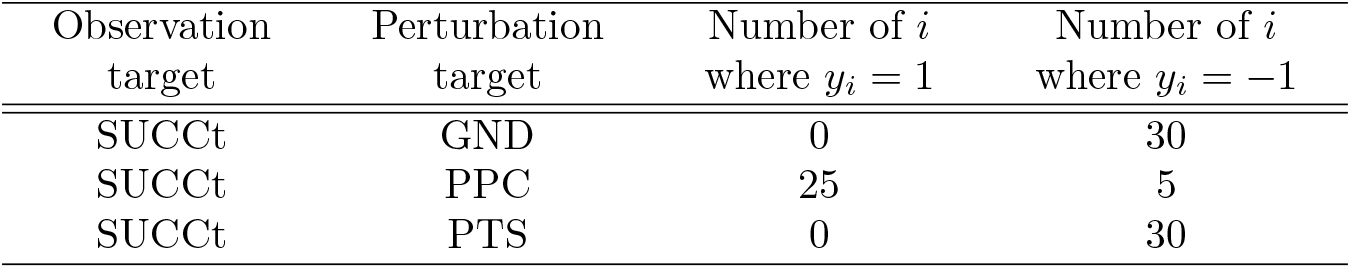
Obtained synthetic perturbation dataset.

#### 3.2.3 Wet lab experiments

We conducted perturbation experiments for the reactions CS, FBP, ICL, LDH, ME1, ME2, PCK, PPC, PPS, and PTA. The genes corresponding to the reactions, listed in Table 2, were introduced into the plasmid vector pLEAD5 (NIPPON GENE CO., LTD.). We amplified the gene sequences using PrimeSTAR GXL DNA Polymerase (Takara Bio Inc.) with *E. coli* DH5*α* (NIPPON GENE CO., LTD.) as a template sequence. The primer sequences were derived from NCBI Genes [45]. The nucleotide sequences of DNAs were analyzed using the BigDye® Terminator v3.1 Cycle Sequencing Kit (Applied Biosystems) with SeqStudio™ Genetic Analyzer (Applied Biosystems). The nucleotide sequence data were processed using GENETYX-Mac NETWORK software, version 15 (GENETYX CORPORATION). We introduced the constructed plasmid vectors into *E. coli* JM109 (NIPPON GENE CO., LTD.). The modified strains were aerobically cultured and anaerobically fermented at 37°C in M9 minimal medium (Thermo Fisher Scientific Inc.). To measure succinate concentrations, we used EnzyChrom Succinate Assay Kit (#ESNT-100, BioAssay Systems) and the absorbance meter SH-8000(CORONA ELECTRIC Co.,Ltd.) at 570 nm. The resulting absorbance, which is a constant multiple of the succinate concentrations, was normalized by the optical density of the bacterial liquid. We calculated the differences between the resulting values and the absorbances of controls, which are of a strain with only pLEAD5, and the signs of the values were used as the real perturbation data. We constructed and measured three replicates for each perturbed reaction. The obtained dataset is shown in Table 2. To validate the overexpression, we performed real-time PCR using QuantStudio5 Real-time PCR system (Thermo Fisher SCIENTIFIC) and confirmed the genes were successfully overexpressed (Supplementary Table S4).

**Table 2:**
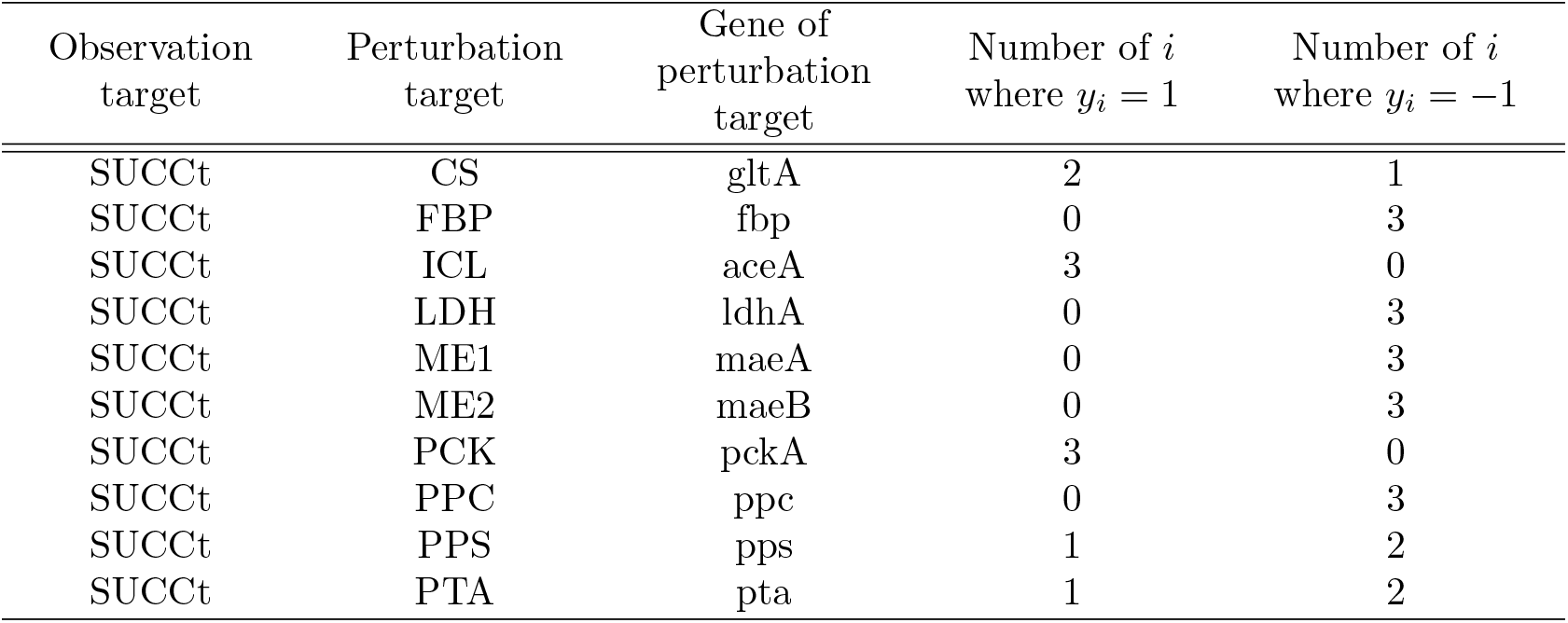
Obtained real perturbation dataset.

### 3.3 Evaluation

#### 3.3.1 Naive Bayes model

To evaluate the effectiveness of incorporating SSA, we constructed a naive Bayes model as follows:

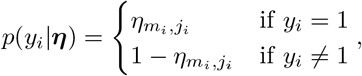

where *η*_*m,j*_ ∈ (0, 1) is a parameter indicating the probability that *y*_*i*_ = 1 where the *i*-th experiment is of (*m, j*), and ***η*** is a matrix whose (*m, j*) element is *η*_*m,j*_. The predictive distribution of one new perturbation record *y*^new^ is as follows:

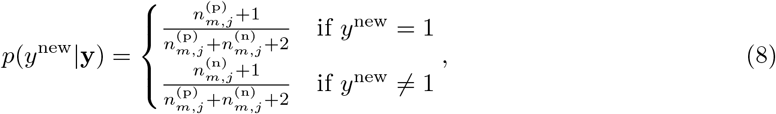

where 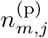 and 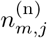 are the numbers of the (*m, j*) experiments in **y** where *y*_*i*_ = 1 and *y*_*i*_ = −1, respectively. We used *ℬ* (*η*_*m,j*_|1, 1) as the prior distribution.

#### 3.3.2 Cross-entropy loss

To evaluate the performance of BayesianSSA, we used the cross-entropy loss as follows:

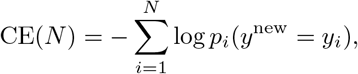

where *N* is the number of trials, which means how many times data are added, and *p*_*i*_(*·*) is the *i*-th predictive distribution of BayesianSSA or the naive Bayes model. The *i*-th predictive distribution is calculated using 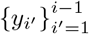. We used *p*_*t*_(*y*^new^ = 1) = 0.5 and *p*_*t*_(*y*^new^ = *y*_*i*_) = *p*_1_(*y*^new^ = *y*_*i*_) as the “random method” and “base method” to calculate the cross-entropy loss, respectively, to be compared with BayesianSSA. Here, *p*_1_(*y*^new^ = *y*_*i*_) indicates the initial distribution of BayesianSSA.

## 4 Results

*m*_name_ denotes the index of the observation target in the following. In other words, the *m*_name_-th observation target is a metabolite or a reaction whose name is “name.” Similarly, *j*_name_ denotes the index of the perturbation target “name” in the following. Unless otherwise stated, we used 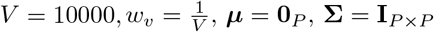, and (*a, b*) = (3, 1) as the hyper-parameters of BayesianSSA where **0**_*P*_ is the *P* –dimensional zero vector, and **I**_*P ×P*_ is the *P × P* identity matrix.

### 4.1 Performance evaluation on synthetic dataset

To compare predictive performance between BayesianSSA and the base method under an ideal situation that the experimental error is low, we examined the cross-entropy loss CE(*N*) (*cf*. the “Crossentropy loss” section) on the synthetic dataset shown in Table 1. Figure 1 shows the cross-entropy loss trajectory for each method. While cross-entropy loss values at the last trial of BayesianSSA with (*a, b*) = (9, 1), (*a, b*) = (6, 2), (*a, b*) = (3, 1), and (*a, b*) = (2, 1) are 19.6, 23.3, 21.4, and 22.3, respectively, those of the random and base methods are 62.4 and 66.6, respectively. BayesianSSA outperformed the random and base method from the perspective of prediction accuracy under the ideal situation.

**Figure 1:**
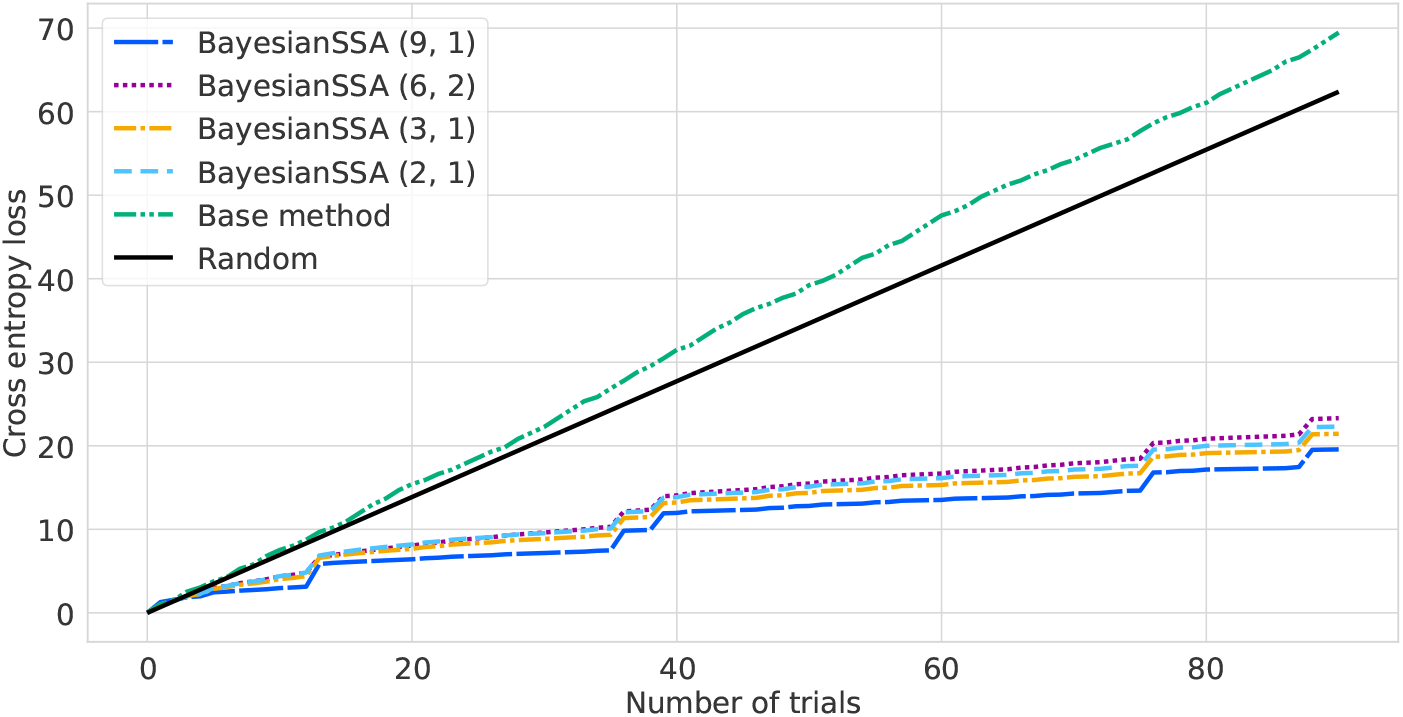
Cross-entropy loss trajectory for each method on the synthetic dataset. The *x*and *y*-axes indicate the number of trials and cross-entropy loss, respectively.

### 4.2 Positivity confidence value transition on pseudo data

To examine the transition of positivity confidence values, we applied BayesianSSA to a pseudo perturbation dataset (shown in Table 3) based on the previous studies [41, 42, 43]. Figure 2 shows the transition of positivity confidence values for increasing SUCCt, *i*.*e*., *c*_*p*_(*m*_SUCCt_, *·*), when fitting BayesianSSA to the pseudo perturbation dataset. We found that the positivity confidence values were updated for multiple reactions rather than one reaction. The first update (from Figures 2(a) to 2(b)) shows the EDA (6PG → G3P + PYR) positivity confidence value for the production of succinate becomes higher. The second update (from Figure 2(b) to Figure 2(c)) shows the reactions included by the flux from FBP to PEP become lower. One of the reaction candidates to increase the SUCCt flux was PPC, whose reaction formula was PEP + CO2 → OAA. While the initial PPC positivity confidence value was 0.714 (Supplementary Table S6), the updated PPC positivity confidence value was 0.869 (Supplementary Table S7). A previous report showed the succinate production of *E. coli* increases when PPC was up-regulated [44], and the PPC positivity confidence value estimated by BayesianSSA is consistent with the report. Similarly, those of GND and PTS were updated from 0.530 and 0.779 to 7.84 *×* 10^−3^ and 3.23 *×* 10^−2^, respectively. Although these results are also consistent with the reports [41, 42, 43], they are naive because the information of these reports was used directly to fit BayesianSSA. The initial positivity confidence values, which were calculated on the basis of the prior distribution, and positivity confidence values updated with the pseudo perturbation dataset are shown in Supplementary Table S6 and S7, respectively.

**Table 3:**
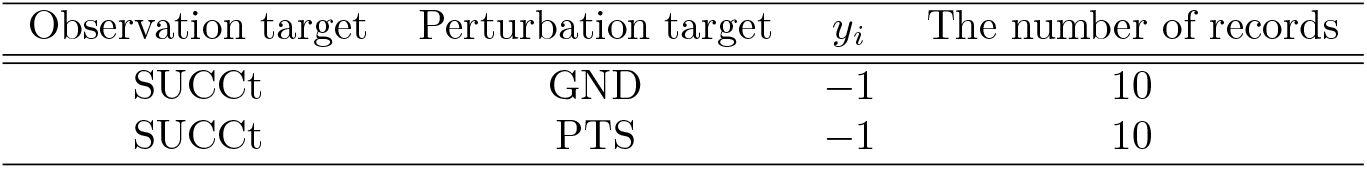
Pseudo perturbation dataset.

**Figure 2:**
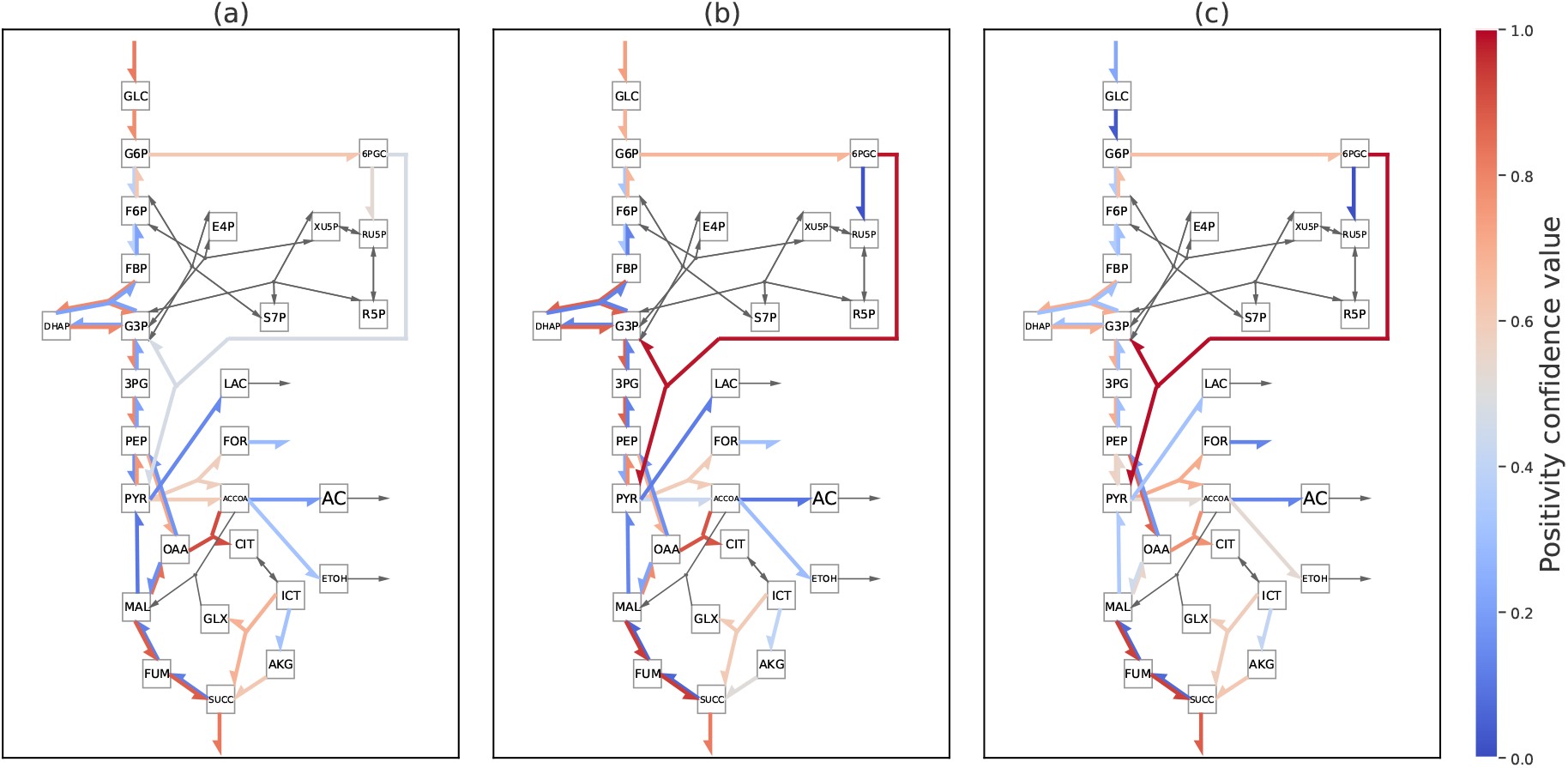
Transition of positivity confidence value for increasing the succinate export flux 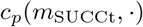. Each square indicates a metabolite in the network. Each arrow indicates a reaction, and its color shows the positivity confidence value (red and blue) or zero response (grey). The reactions corresponding to the edges in this figure are shown in Supplementary Table S5. **(a)** Values given by the initial model. **(b)** Values given by the model updated by 10 perturbation records that *y*_*i*_ = −1 where *m*_*i*_ = *m*_SUCCt_ and *j*_*i*_ = *j*_GND_. **(c)** Values given by the model updated by 10 perturbation records that *y*_*i*_ = −1 where *m*_*i*_ = *m*_SUCCt_ and *j*_*i*_ = *j*_PTS_ in addition to the perturbation records of **(b)**.

### 4.3 Performance evaluation on real data

To compare the performances between BayesianSSA and the naive Bayes model, which uses only data without SSA, we examined the cross-entropy loss CE(*N*) (*cf*. the “Cross-entropy loss” section). Figure 3 shows the cross-entropy loss trajectory for each method. The perturbation records were randomly shuffled and used in each trial. The loss of BayesianSSA is comparable to that of the base method in the early trials but smaller in the late trials. While the cross-entropy loss values at the last trial of BayesianSSA with (*a, b*) = (9, 1), (*a, b*) = (6, 2), (*a, b*) = (3, 1), and (*a, b*) = (2, 1) are 14.8, 15.7, 15.4, and 16.1, respectively, those of the random method, the base method, and the naive Bayes model are 20.8, 19.0, and 17.2, respectively. BayesianSSA outperformed the random and base methods and the naive Bayes model from the perspective of prediction accuracy. While BayesianSSA made a prediction on the basis of the perturbation dataset and SSA, the prediction of the base method is only based on SSA. This result suggests that BayesianSSA can integrate environmental information of the real dataset into SSA predictions. Similarly, the difference between BayesianSSA and the naive Bayes is whether or not SSA is incorporated. This result also indicates the practicability of BayesianSSA and that incorporating SSA into statistical models improves predictive performance.

**Figure 3:**
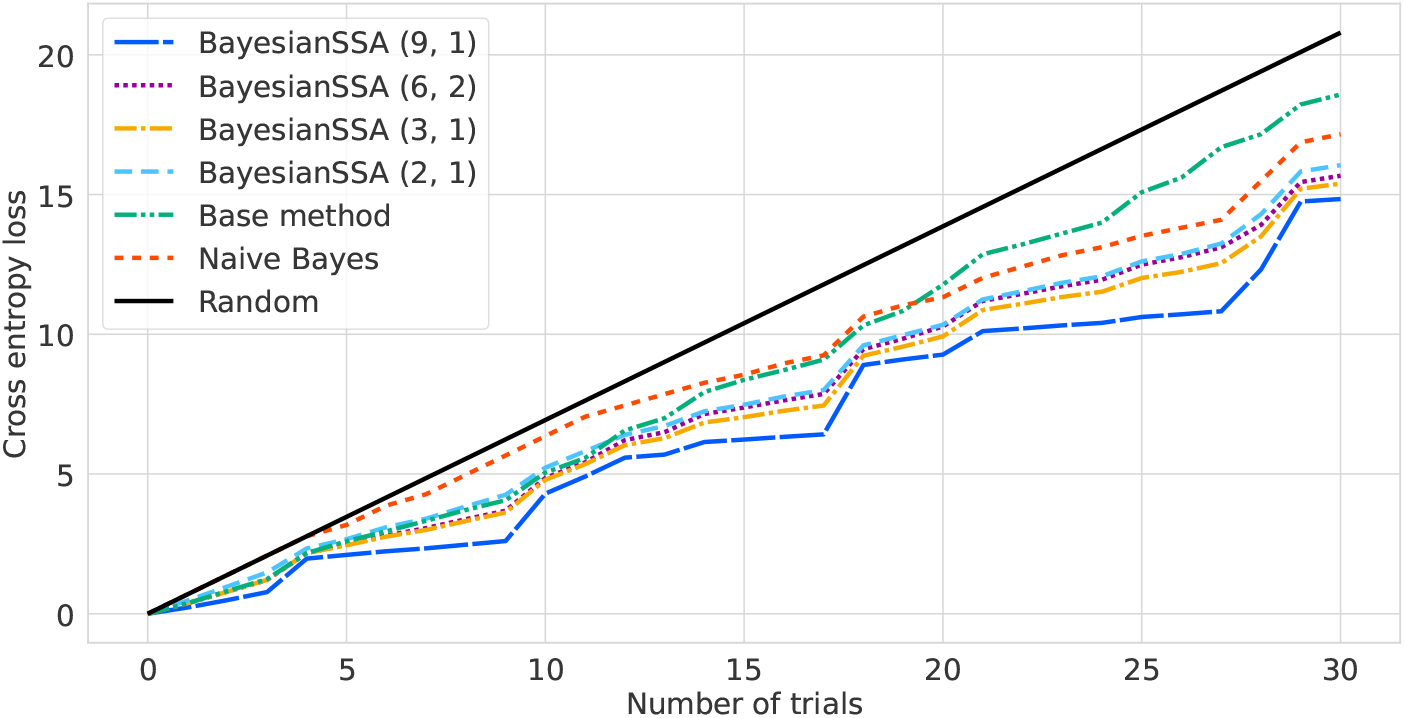
Cross-entropy loss trajectory for each method on the real dataset. The *x*and *y*-axes indicate the number of trials and cross-entropy loss, respectively.

The main difference between BayesianSSA and the naive Bayes model is in the predictions for out-of-sample perturbations, which are of new (*m, j*) experiments. To calculate the predictive distribution of a (*m, j*) experiment, the naive Bayes model uses only the results of (*m, j*) experiments (Eq. (8)) while BayesianSSA uses the results of all the (*m, j*) ∈ Λ experiments (Eq. (6)). That is, BayesianSSA can leverage data to predict the responses to out-of-sample perturbations through the **r** posterior distribution. To validate the predictive performance for out-of-sample perturbations, we examined the predictive probabilities by splitting the real dataset. For example, the predictive probabilities of the three replicates for the reaction CS were calculated by fitting BayesianSSA to the real dataset except for CS. Figure 4 shows the distribution of predictive probabilities calculated by BayesianSSA for out-of-sample perturbations. Here, each replicate was evaluated separately. The median of the distribution was 0.67, which is better than random (0.5), and this result indicates that fitting BayesianSSA contributes to the predictive performance for out-of-sample perturbations.

**Figure 4:**
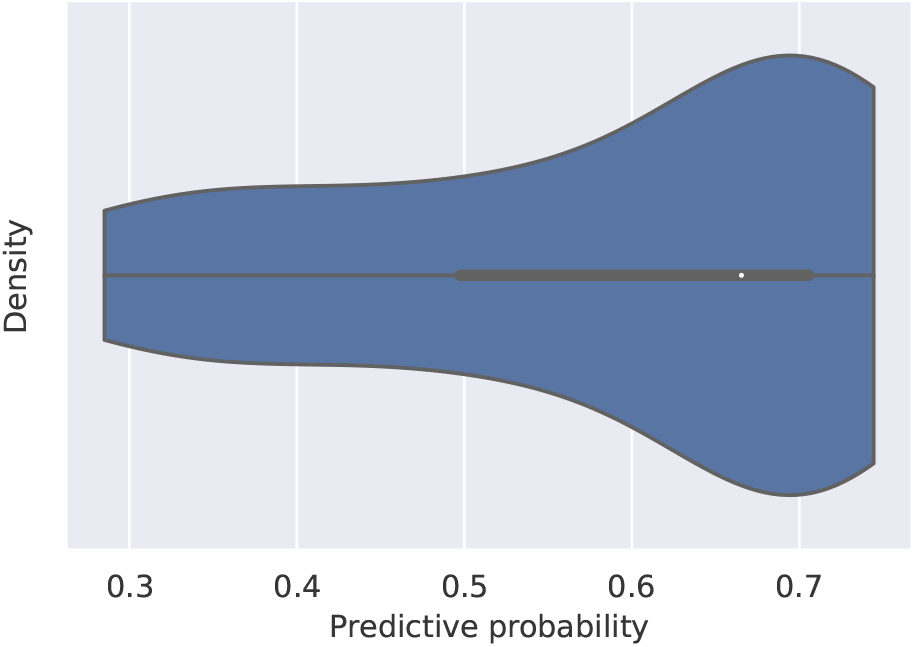
Distribution of predictive probabilities calculated by BayesianSSA for out-of-sample perturbations. Each value was calculated for the real dataset for new (*m, j*) perturbation. The white dot indicates the median of the distribution.

### 4.4 Positivity confidence values on real data

To interpret the BayesianSSA estimation results, we examined the positivity confidence values after BayesianSSA was fitted to the real data whose observation target is the succinate export flux. Figure 5 shows the positivity confidence values in the metabolic network. All positivity confidence values updated by the real perturbation dataset are shown in Supplementary Table S8. We found high positivity confidence values of THD r and NDH (*c*_*p*_(*m*_SUCCt_, *j*_THD r_) = 0.95, *c*_*p*_(*m*_SUCCt_, *j*_NDH_) = 0.92), which both use NADH. NDH produces Q8H2, which is required for FRD. In the tricarboxylic acid (TCA) cycle, ICL, MDH, FRD, FUM r, and CS show high positivity confidence values (*>* 0.85). We also applied BayesianSSA with another prior distribution of **r**, which is based on log-normal distributions with random parameters, to the real dataset (Supplementary Figure S1). All positivity confidence values using this prior distribution are shown in Supplementary Table S9. The reactions with *c*_*p*_(*m, j*) *>* 0.85 are almost shared in the two BayesianSSA results. The only difference is the presence or absence of NDH (*cf*. Supplementary Table S8 and Supplementary Table S9). These results suggest that BayesianSSA is robust to changes in the prior distributions of **r**. We have also examined other observation targets besides succinate export flux and found that they can also be updated to high positivity confidence values (Supplementary Figure S2 and S3).

**Figure 5:**
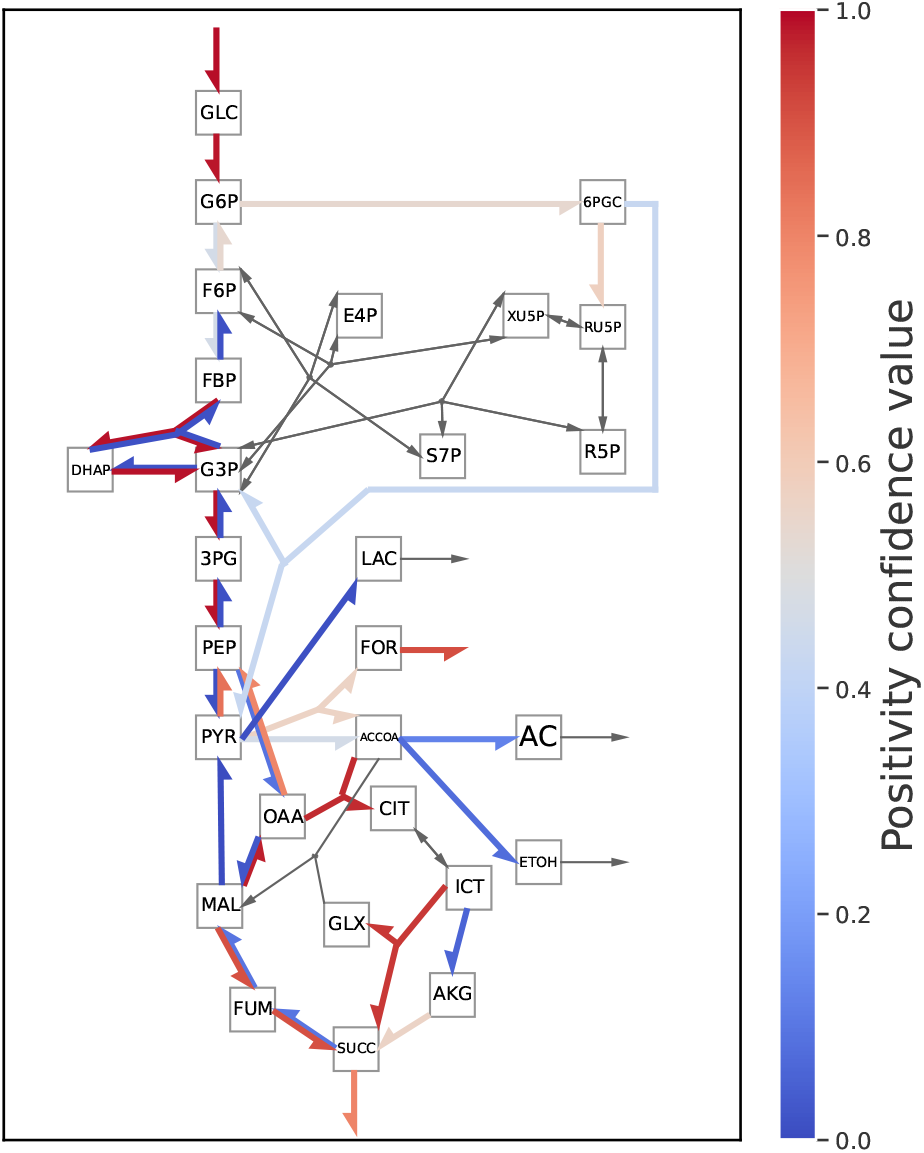
Positivity confidence values by BayesianSSA to increase the succinate export flux on metabolic networks. The positivity confidence values were calculated by BayesianSSA fitted to real data. Each square indicates a metabolite in the network. Each arrow indicates a reaction, and its color shows the positivity confidence value (red and blue) or zero response (grey). The reactions corresponding to the edges in this figure are shown in Supplementary Table S5.

Positivity confidence values calculated by the BayesianSSA posterior distribution were consistent with previous reports. The positivity confidence values of CS and ICL, which are included in the glyoxylate pathway, were high. The glyoxylate pathway was reported as an essential pathway for succinate production [46, 47]. Similarly, the reductive pathway, which includes FUM r and FRD, was reported as another essential pathway [46, 47]. Despite these consistencies, other previous reports for several reactions, such as PPC [44] and PTS [42, 43] are inconsistent with our results.

### 4.5 Posterior distribution of r

To examine the differences between the prior and posterior distribution of **r**, we compared *p*(**r**|**y**) with *p*(**r**). Figure 6 shows *p*(**r**) and *p*(**r**|**y**) on the first and second principal components of 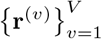. We found that the posterior distribution had several peaks where the prior distribution only had one peak.

**Figure 6:**
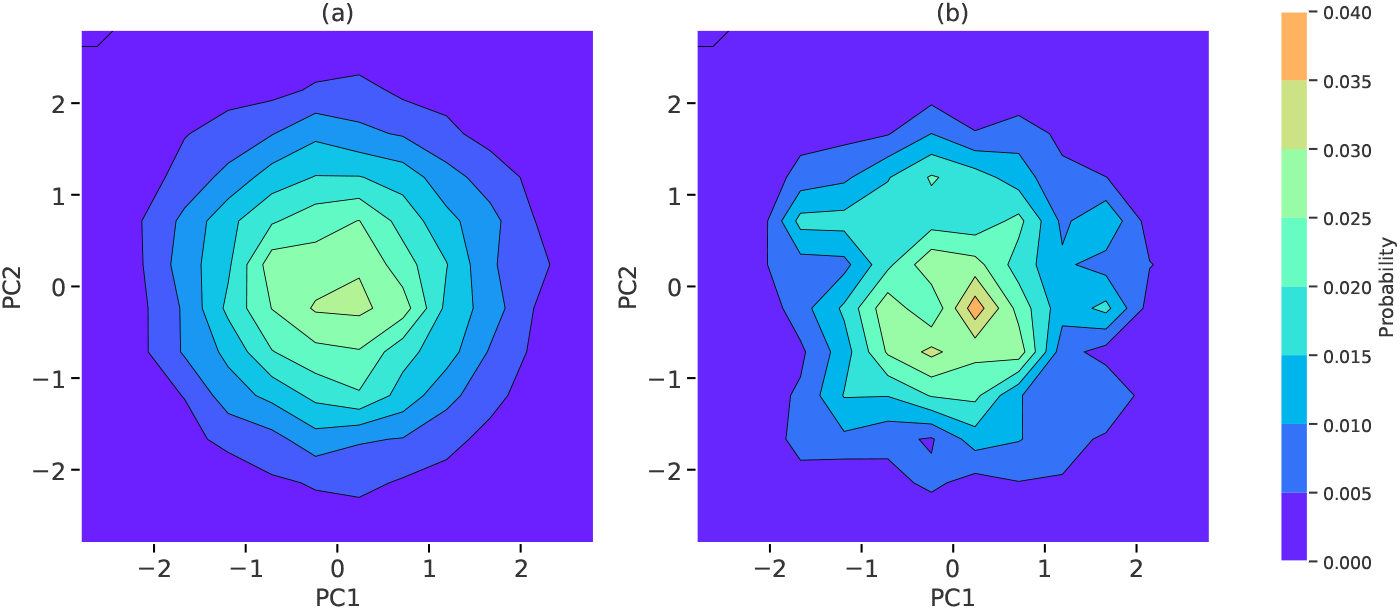
Contour plots of the prior **(a)** and posterior **(b)** distributions of log **r**. The *x*and *y*-axes indicate the first and second principal components, respectively.

This result indicates that the distribution of **r** is tailored to several cases of environments in which the used dataset was collected.

## 5 Discussion

We demonstrated the effectiveness of BayesianSSA on the basis of predictive performance using the real perturbation dataset where the observation target is the succinate export flux (SUCCt). BayesianSSA outperformed the base method (Figure 3), which is not fitted to the dataset, and this result showed BayesianSSA could integrate environmental information into SSA predictions. For validation of practicability, BayesianSSA also outperformed the random method and the naive Bayes model, and the performance of BayesianSSA for out-of-sample perturbations, which are of new (*m, j*) experiments, was better than random. These results show that BayesianSSA can consider the relationships between different (*m, j*) perturbations through **r**, and that this consideration contributes to predictive performance.

We considered *ρ*_*m,j*_ as a parameter and set the prior distribution in this study. As another option, *ρ*_*m,j*_ can be given by an error rate of the measurement equipment used for the perturbation experiment. For example, consider a case where the error distribution of the used measurement equipment is a normal distribution with a mean parameter that equals the true value and a variance parameter *σ*^2^ = 1 as an error distribution. If the experimental value obtained by the perturbation is 1, the probability that the true value is less than zero is approximately 16%. Therefore, setting 1 − *ρ*_*m,j*_ = 0.16 can make BayesianSSA consider the error distribution of the measurement equipment. In this way, we can set a certain value of *ρ*_*m,j*_. Note that we can easily calculate the posterior distribution of this model (Supplementary Section S4).

There are two directions for future work related to **r**. First, the updated *p*(**r**) may be used for response predictions in another metabolic network. When the reaction rate function *F*_*j*_ and the probability distribution of **x** are equal between the two metabolic networks, *p*(*r*_*j,m*_) can be used in another system that includes the *i*-th reaction and the *m*-th metabolite. Second, as previously discussed [10], we can consider allosteric regulation. Allosteric regulation is a type of regulation that increases/decreases reaction rates as a metabolite concentration increases [48, 49]. We can easily consider allosteric regulation by setting *r*_*j,m*_ ≠ 0. Technically, *r*_*j,m*_ *<* 0 can be implemented by reversing the sign of *r*_*j,m*_ after sampling **r**^(*v*)^. However, we need to know the (*j, m*) pairs that have allosteric regulation in advance, and we omitted considering allosteric regulations in this study.

There is room for choice regarding the prior distribution of **r**. First, continuous distributions can be adopted. We used the empirical distribution with samples from log-normal distributions as the **r** prior distribution for all experiments (*cf*. Eq. (4)). As long as the constraint on **r** is satisfied, other distributions can be adopted. However, the likelihood function changes discretely (*cf*. Eq. (3)), and the advantage of adopting a continuous distribution with employing MCMC methods may be limited.

Second, ensemble approaches for several types of prior distributions may be effective. As shown in Figure 5 and Supplementary Figure S1, the effect of the prior distribution is not negligible when dealing with a limited sample size. Using several types of prior distributions may contribute to making robust predictions.

Although we omitted the biomass production processes in this study, considering them can improve the representation of real biological activity. One commonly used approach in FBA involves optimizing the biomass objective function [18, 19]. However, since the biomass objective function depends on the specific strains and environmental conditions [50], it is difficult to use the biomass objective function when analyzing unfamiliar strains. Another difficulty is that the biomass objective function is a pseudo-reaction that contains multiple reactions and cannot be treated kinetically. Unlike FBA-based methods, which can consider biomass production processes as a single reaction, kinetics-based methods including BayesianSSA need to faithfully model the biomass production process. That is, it is necessary to define the biomass production rate equations as in a previous study [21].

Utilizing the positivity confidence value calculation (the “Positivity confidence value” section) and Bayesian updating (the “Bayesian updating” section) in BayesianSSA, we can construct an iterative design-build-test-learn (DBTL) cycle [51] on the basis of BayesianSSA for proposals of reactions to be perturbed. Specifically, the procedure is as follows:

1. Calculate positivity confidence values by Eq. (7).
2. Obtain a proposal of which perturbation and observation targets are validated in accordance with positivity confidence values.
3. Conduct perturbation experiments in accordance with the proposal.
4. Update the posterior distributions by Eq. (5).
5. Return to the first step.

One advantage of this scheme is the high efficiency because the experimental validation of proposals obtained by BayesianSSA is also a process collecting data for updating the BayesianSSA posterior distribution.

## 6 Conclusions

In this study, we proposed BayesianSSA, a Bayesian statistical model based on SSA. SSA was previously developed as a method to predict qualitative responses to enzyme perturbations on the basis of the structural information of the reaction network. However, the network structural information can sometimes be insufficient to predict qualitative responses unambiguously, which is a practical issue in bioproduction applications. To address this, BayesianSSA extracts environmental information from perturbation datasets collected in environments of interest and integrates it into SSA predictions. We applied BayesianSSA to synthetic and real datasets of the central metabolic pathway of *E. coli*. As a result, BayesianSSA outperformed the base method, which is the same as the BayesianSSA model but utilizes an initial prior distribution without incorporating perturbation datasets. This result shows that BayesianSSA can successfully integrate environmental information extracted from perturbation data into SSA predictions. In addition, the positivity confidence values estimated by BayesianSSA for increasing the succinate export flux were consistent with the known pathways reported to enhance the flux in previous studies. We believe that BayesianSSA will accelerate the chemical bioproduction process and contribute to advancements in the field.

## Supporting information

Supplementary data including Figures S1-3, Tables S1-S9, and Sections S1-S4.

## Declarations

### Ethics approval and consent to participate

Not applicable.

### Consent for publication

Not applicable.

### Availability of data and materials

Supplementary material is available from the journal website.

The implementation of the algorithm is available on GitHub (https://github.com/shion-hosoda-hitachi/BayesianSSA).

### Authors’ contributions

S.H., T.O., A.M., and M.S. conceptualized this study. S.H. devised the model, designed the algorithms, implemented the software, and performed all the computational experiments. H.I. performed all the wet lab experiments. T.M., H.I., and M.T. established the wet lab experimental procedures. S.H., M.S., T.O., and A.M. interpreted the computational results. S.H. and M.S. investigated previous researches for biological insights. S.H. wrote the draft. T.O., A.M., M.S., M.T., H.I., and T.M. revised the manuscript critically. All authors read and approved the final manuscript.

## Acknowledgements

We would like to thank Y. Mizunuma for her technical assistance with the wet lab experiments. We also thank Dr. K. Yokoyama for technical guidance. We appreciate the advice and technical support with the wet lab experiments from Dr. T. Takeya. We are grateful to Dr. A. Kandori, Dr. K. Watanabe, and Dr. S. Yabuuchi for their guidance throughout our project. This research did not receive any specific grant from funding agencies in the public, commercial, or not-for-profit sectors.

