## Supplementary data including Figures S1-3, Tables S1-S9, and Sections S1-S4. for "BayesianSSA: a Bayesian statistical model based on structural sensitivity analysis for predicting responses to enzyme perturbations in metabolic networks"

#### S1 Posterior distribution

$$
\begin{aligned}
p(\mathbf{r}=\mathbf{r}^{(v)},\boldsymbol{\rho}|\mathbf{y}) &\propto p(\mathbf{y}|\mathbf{r}=\mathbf{r}^{(v)},\boldsymbol{\rho})p(\mathbf{r}=\mathbf{r}^{(v)})p(\boldsymbol{\rho}) \\
&= \left(\prod_{m=1}^{M+J}\prod_{j=1}^{J+L}\rho_{m,j}^{t_{m,j,v}}(1-\rho_{m,j})^{f_{m,j,v}}\right)w_v \\
&\qquad \left(\prod_{m=1}^{M+J}\prod_{j=1}^{J+L}\frac{\rho_{m,j}^{a-1}(1-\rho_{m,j})^{b-1}}{\mathrm{Beta}(a,b)}\right) \\
&\propto w_v\prod_{m=1}^{M+J}\prod_{j=1}^{J+L}\rho_{m,j}^{\hat{a}_{m,j,v}-1}(1-\rho_{m,j})^{\hat{b}_{m,j,v}-1}.
\end{aligned}
$$

$$
p(\mathbf{r}=\mathbf{r}^{(v)},\boldsymbol{\rho}|\mathbf{y}) = \frac{w_v\prod_{m=1}^{M+J}\prod_{j=1}^{J+L}\rho_{m,j}^{\hat{a}_{m,j,v}-1}(1-\rho_{m,j})^{\hat{b}_{m,j,v}-1}}{\sum_{v'=1}^{V}w_{v'}\prod_{m=1}^{M+J}\prod_{j=1}^{J+L}\mathrm{Beta}\left(\hat{a}_{m,j,v'},\hat{b}_{m,j,v'}\right)}.
$$

#### S2 Predictive distribution

$$\begin{aligned}
p(\mathbf{y}^{\text{new}}|\mathbf{y}) &= \sum_{v=1}^{V} \int p(\mathbf{y}^{\text{new}}|\mathbf{r}=\mathbf{r}^{(v)}, \boldsymbol{\rho}) p(\mathbf{r}=\mathbf{r}^{(v)}, \boldsymbol{\rho}|\mathbf{y}) d\boldsymbol{\rho} \\
&= \sum_{v=1}^{V} \int \left( \prod_{m=1}^{M+J} \prod_{j=1}^{J+L} \rho_{m,j}^{t^{\text{new}}_{m,j,v}} (1-\rho_{m,j})^{f^{\text{new}}_{m,j,v}} \right) \\
&\qquad \frac{w_v \prod_{m=1}^{M+J} \prod_{j=1}^{J+L} \rho_{m,j}^{\hat{a}_{m,j,v}-1} (1-\rho_{m,j})^{\hat{b}_{m,j,v}-1}}{\sum_{v'=1}^{V} w_{v'} \prod_{m=1}^{M+J} \prod_{j=1}^{J+L} \text{Beta}\Big(\hat{a}_{m,j,v'}, \hat{b}_{m,j,v'}\Big)} d\boldsymbol{\rho} \\
&= \sum_{v=1}^{V} \int \frac{w_v \prod_{m=1}^{M+J} \prod_{j=1}^{J+L} \rho_{m,j}^{\hat{a}_{m,j,v}+t^{\text{new}}_{m,j,v}-1} (1-\rho_{m,j})^{\hat{b}_{m,j,v}+f^{\text{new}}_{m,j,v}-1}}{\sum_{v'=1}^{V} w_{v'} \prod_{m=1}^{M+J} \prod_{j=1}^{J+L} \text{Beta}\Big(\hat{a}_{m,j,v'}, \hat{b}_{m,j,v'}\Big)} d\boldsymbol{\rho} \\
&= \frac{\sum_{v=1}^{V} w_v \prod_{m=1}^{M+J} \prod_{j=1}^{J+L} \text{Beta}\Big(\hat{a}_{m,j,v}+t^{\text{new}}_{m,j,v}, \hat{b}_{m,j,v}+f^{\text{new}}_{m,j,v}\Big)}{\sum_{v'=1}^{V} w_{v'} \prod_{m=1}^{M+J} \prod_{j=1}^{J+L} \text{Beta}\Big(\hat{a}_{m,j,v'}, \hat{b}_{m,j,v'}\Big)} \\
&= \frac{\sum_{v=1}^{V} w_v \prod_{(m,j)\in\Lambda^{\text{new}}} \text{Beta}\Big(\hat{a}_{m,j,v}+t^{\text{new}}_{m,j,v}, \hat{b}_{m,j,v}+f^{\text{new}}_{m,j,v}\Big)}{\sum_{v'=1}^{V} w_{v'} \prod_{(m,j)\in\Lambda} \text{Beta}\Big(\hat{a}_{m,j,v'}, \hat{b}_{m,j,v'}\Big)}.
\end{aligned}$$

#### S3 Bayesian update

The updated posterior distribution $p(\mathbf{r}|\mathbf{y}^{\text{new}}, \mathbf{y})$ can be written by the former posterior distribution $p(\mathbf{r}|\mathbf{y})$.

$$\begin{aligned}
p(\mathbf{r}=\mathbf{r}^{(v)}|\mathbf{y}^{\text{new}}, \mathbf{y}) &= \int p(\mathbf{r}=\mathbf{r}^{(v)}, \boldsymbol{\rho}|\mathbf{y}^{\text{new}}, \mathbf{y}) d\boldsymbol{\rho} \\
&= \int \frac{p(\mathbf{y}^{\text{new}}|\mathbf{r}=\mathbf{r}^{(v)}, \boldsymbol{\rho}) p(\mathbf{r}=\mathbf{r}^{(v)}, \boldsymbol{\rho}|\mathbf{y})}{p(\mathbf{y}^{\text{new}}|\mathbf{y})} d\boldsymbol{\rho} \\
&\propto \int p(\mathbf{y}^{\text{new}}|\mathbf{r}=\mathbf{r}^{(v)}, \boldsymbol{\rho}) p(\mathbf{r}=\mathbf{r}^{(v)}, \boldsymbol{\rho}|\mathbf{y}) d\boldsymbol{\rho} \\
&= \int \left( \prod_{i=1}^{N} \rho_{m_i,j_i}^{\mathbb{I}_{y_i^{\text{new}}=q_{m_i,j_i}(\mathbf{r})}} (1-\rho_{m_i,j_i})^{\mathbb{I}_{y_i^{\text{new}}\neq q_{m_i,j_i}(\mathbf{r})}} \right) \\
&\qquad \frac{w_v \prod_{m=1}^{M+J} \prod_{j=1}^{J+L} \rho_{m,j}^{\hat{a}_{m,j,v}-1} (1-\rho_{m,j})^{\hat{b}_{m,j,v}-1}}{\sum_{v'=1}^{V} w_{v'} \prod_{m=1}^{M+J} \prod_{j=1}^{J+L} \text{Beta}\Big(\hat{a}_{m,j,v'}, \hat{b}_{m,j,v'}\Big)} d\boldsymbol{\rho} \\
&\propto w_v \int \prod_{m=1}^{M+J} \prod_{j=1}^{J+L} \rho_{m,j}^{\hat{a}_{m,j,v}+t^{\text{new}}_{m,j,v}-1} (1-\rho_{m,j})^{\hat{b}_{m,j,v}+f^{\text{new}}_{m,j,v}-1} d\boldsymbol{\rho} \\
&= w_v \prod_{m=1}^{M+J} \prod_{j=1}^{J+L} \text{Beta}\Big(\hat{a}_{m,j,v}+t^{\text{new}}_{m,j,v}, \hat{b}_{m,j,v}+f^{\text{new}}_{m,j,v}\Big).
\end{aligned}$$

Thus,

$$p(\mathbf{r}=\mathbf{r}^{(v)}|\mathbf{y}^{\text{new}}, \mathbf{y}) = \frac{h(\mathbf{y}, \mathbf{r}^{(v)})}{\sum_{v'=1}^{V} h(\mathbf{y}, \mathbf{r}^{(v')})},$$

where

$$h(\mathbf{y}, \mathbf{r}^{(v)}) = w_v \prod_{(m,j)\in\Lambda} \mathrm{Beta}\Big(\hat{a}_{m,j,v} + t^{\mathrm{new}}_{m,j,v}, \hat{b}_{m,j,v} + f^{\mathrm{new}}_{m,j,v}\Big),$$
$$\Lambda = \big\{(m, j) \mid t_{m,j,v} + f_{m,j,v} + t^{\mathrm{new}}_{m,j,v} + f^{\mathrm{new}}_{m,j,v} \neq 0\big\}.$$

### S4 Case that $\boldsymbol{\rho}$ is hyper-parameter

The posterior distribution in BayesianSSA can be calculated as follows:

$$p(\mathbf{r} = \mathbf{r}^{(v)}|\mathbf{y}) = \frac{f(\mathbf{y}, \mathbf{r}^{(v)})}{\sum_{v'} f(\mathbf{y}, \mathbf{r}^{(v')})},$$

where

$$\begin{aligned} f(\mathbf{y}, \mathbf{r}^{(v)}) &= p(\mathbf{y}|\mathbf{r} = \mathbf{r}^{(v)})p(\mathbf{r} = \mathbf{r}^{(v)}) \\ &= \left(\prod_{i\in T} \rho_i\right)\left(\prod_{i\in F}(1 - \rho_i)\right) w_v, \\ T &= \{i \mid y_i = q_{m_i,j_i}(\mathbf{r})\}, \\ F &= \{i \mid y_i \neq q_{m_i,j_i}(\mathbf{r})\}. \end{aligned}$$

We can calculate the posterior distribution using the product of the values of the prior and likelihood distribution and their normalization. We implemented these calculation using the log-sum-exp trick.

### S5 Figures and Tables

Table S1: Example of SSA application results.

| Prediction | Count |
|---|---|
| Positive | 0 |
| Negative | 0 |
| Zero | 24 |
| Indefinite | 56 |

Each number shows a count of SSA predictive responses (positive, negative, zero, or indefinite) of the succinate export flux to perturbations of reactions in the central metabolic pathway of *Escherichia coli* (shown in the “Metabolic network information” section in the main manuscript).

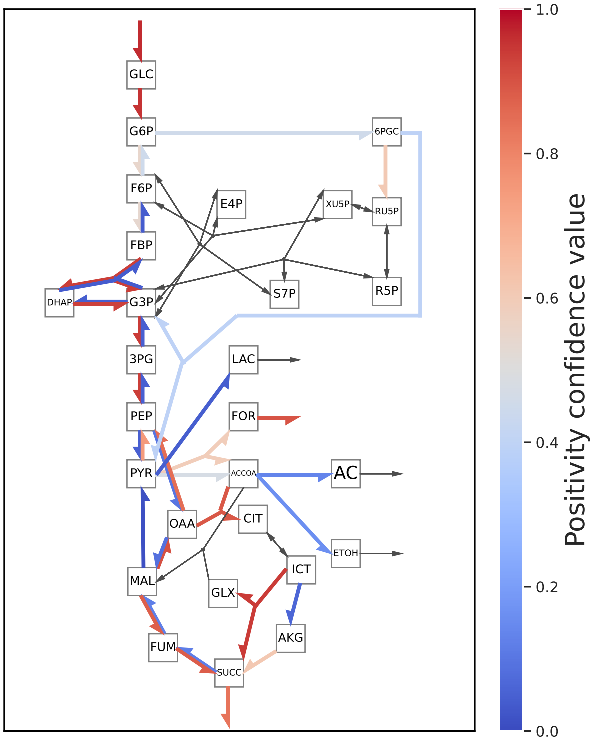


Figure S1: Positivity confidence values mapped on the metabolic network. The positivity confidence values were calculated by BayesianSSA fitted to real data using log-normal distributions with random parameters to increase the succinate export flux. Each square indicates a metabolite in the network. Each arrow indicates a reaction, and its color shows the positivity confidence value (red and blue) or zero response (grey). The reactions corresponding to the edges in this figure are shown in Supplementary Table S5.

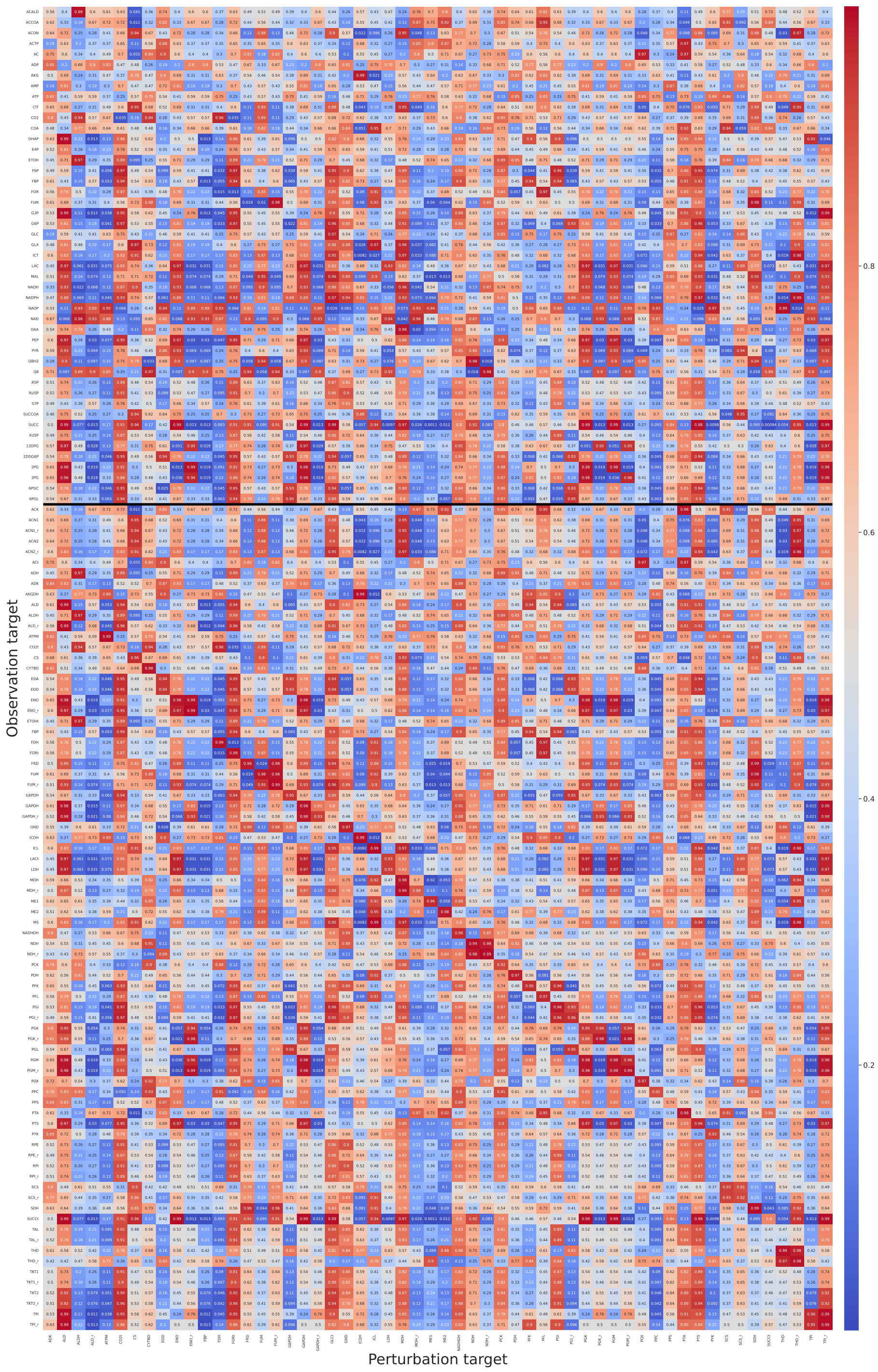


Figure S2: Positivity confidence value matrix. The positivity confidence values were calculated by BayesianSSA fitted to real data. The $x$-axis indicates the perturbation target. The $y$-axis indicates the observation target, the affected reaction flux (below the black line) or metabolite concentration (above the black line). Each element shows the positivity confidence value (red and blue). Positive, negative, and zero responses were omitted.

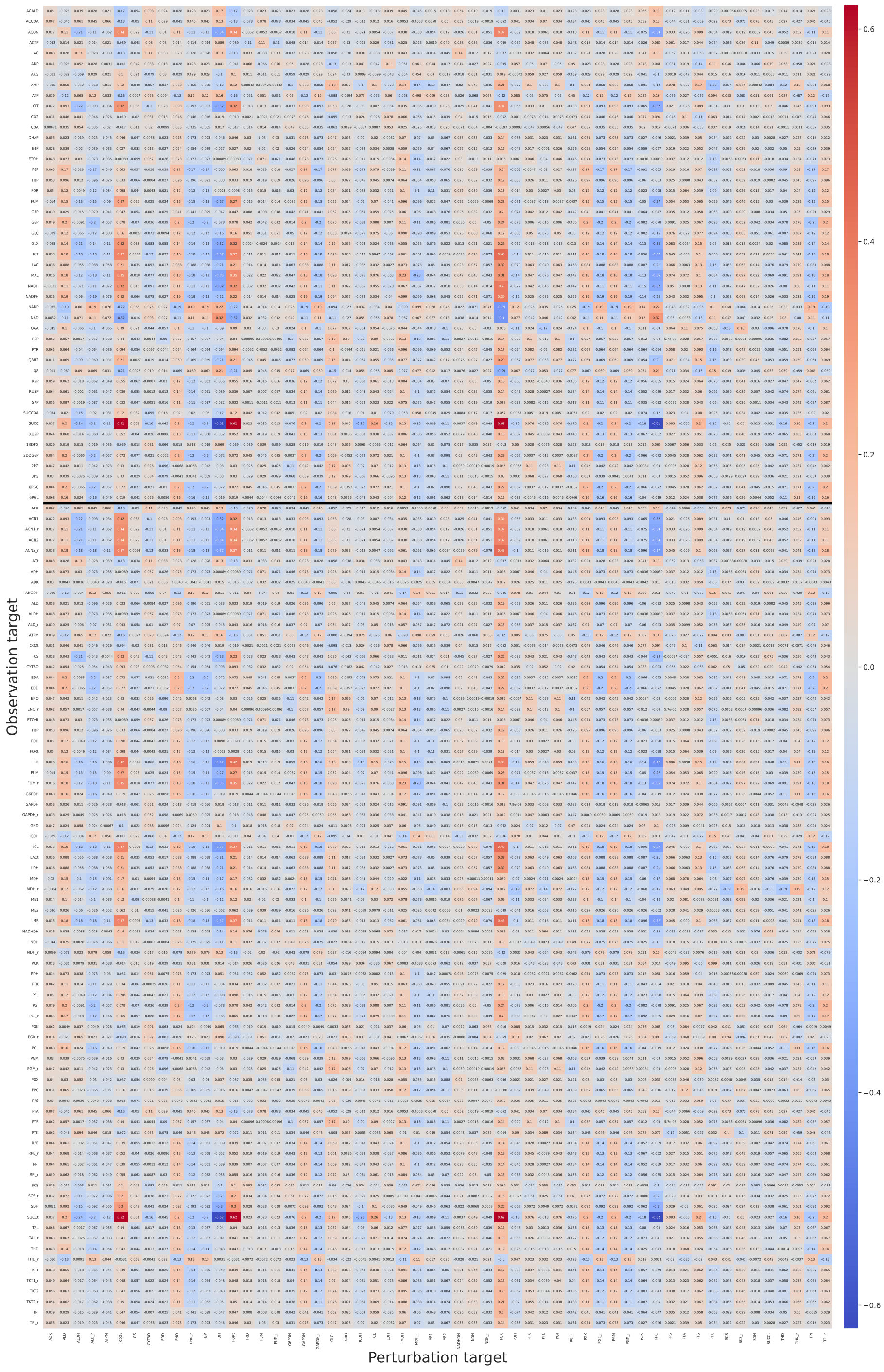


Figure S3: Update difference of positivity confidence value matrix. The positivity confidence values were calculated by BayesianSSA fitted to real data and BayesianSSA with the prior distribution. The $x$-axis indicates the perturbation target. The $y$-axis indicates the observation target, the affected reaction flux (below the black line) or metabolite concentration (above the black line). Each element shows the update difference of positivity confidence value (red and blue). Positive, negative, and zero responses were omitted.

Table S2: Metabolite abbreviations

| Abbreviation | Name |
|---|---|
| AC | Acetate |
| ACALD | Acetaldehyde |
| ACCOA | Acetyl-CoA |
| ACON | Cis-aconitate |
| ACTP | Acetyl phosphate |
| AKG | $\alpha$-ketoglutarate |
| CIT | Citrate |
| DHAP | Dihydroxyacetonephosphate |
| E4P | Erythrose-4-phosphate |
| ETOH | Ethanol |
| F6P | Fructose-6-phosphate |
| FBP | Fructose-1,6-bisphosphate |
| FOR | Formate |
| FUM | Fumarate |
| G3P | Glyceraldehyde-3-phosphate |
| G6P | Glucose-6-phosphate |
| GLC | Glucose |
| GLX | Glyoxylate |
| ICT | Isocitrate |
| LAC | Lactate |
| MAL | Malate |
| OAA | Oxaloacetate |
| PEP | Phosphoenolpyruvate |
| PYR | Pyruvate |
| Q8 | Quinone |
| Q8H2 | Quinol |
| R5P | Ribose-5-phosphate |
| RU5P | Ribulose-5-phosphate |
| S7P | Sedoheptulose-7-phosphate |
| SUCC | Succinate |
| SUCCOA | Succinyl-CoA |
| XU5P | Xylulose-5-phosphate |
| 13DPG | 1,3-diphosphoglycerate |
| 2DDG6P | 2-dehydro-3-deoxy-gluconate-6-phosphate |
| 2PG | 2-phosphoglycerate |
| 3PG | 3-phosphoglycerate |
| 6PGC | 6-phosphogluconate |
| 6PGL | 6-phosphogluconolacton |

Table S3: Reaction names and corresponding equations

| Reaction name | Equation |
|---|---|
| ACK | ACTP + ADP → AC + ATP |
| ACN1 | CIT → ACON |
| ACN1_r | ACON → CIT |
| ACN2 | ACON → ICT |
| ACN2_r | ICT → ACON |
| ACt | AC → (Export) |
| ADH | ACALD + NADH → ETOH + NAD |
| ADK | AMP + ATP → 2 ADP |
| AKGDH | AKG + COA + NAD → CO2 + NADH + SUCCOA |
| ALD | FBP → DHAP + G3P |
| ALDH | ACCOA + NADH → ACALD + COA + NAD |
| ALD_r | DHAP + G3P → FBP |
| ATPM | ATP → ADP |
| CO2t | CO2 → (Export) |
| CS | ACCOA + OAA → CIT + COA |
| CYTBO | ADP + Q8H2 → ATP + Q8 |
| EDA | 2DDG6P → G3P + PYR |
| EDD | 6PGC → 2DDG6P |
| ENO | 2PG → PEP |
| ENO_r | PEP → 2PG |
| ETOHt | ETOH → (Export) |
| FBP | FBP → F6P |
| FDH | FOR → CO2 |
| FORt | FOR → (Export) |
| FRD | FUM + Q8H2 → Q8 + SUCC |
| FUM | FUM → MAL |
| FUM_r | MAL → FUM |
| G6PDH | G6P + NADP → NADPH + 6PGL |
| GAPDH | G3P + NAD → NADH + 13DPG |
| GAPDH_r | NADH + 13DPG → G3P + NAD |
| GLCt | (Import) → GLC |
| GLK | ATP + GLC → ADP + G6P |
| GND | NADP + 6PGC → CO2 + NADPH + RU5P |
| ICDH | ICT + NADP → AKG + CO2 + NADPH |
| ICL | ICT → GLX + SUCC |
| LACt | LAC → (Export) |
| LDH | NADH + PYR → LAC + NAD |
| MDH | MAL + NAD → NADH + OAA |
| MDH_r | NADH + OAA → MAL + NAD |
| ME1 | MAL + NAD → CO2 + NADH + PYR |
| ME2 | MAL + NADP → CO2 + NADPH + PYR |
| MS | ACCOA + GLX → COA + MAL |
| NADHDH | 2 ADP + NADH → 2 ATP + NAD |
| NDH | NADH + Q8 → NAD + Q8H2 |
| NDH_r | NAD + Q8H2 → NADH + Q8 |
| PCK | ATP + OAA → ADP + CO2 + PEP |
| PDH | COA + NAD + PYR → ACCOA + CO2 + NADH |

Continued on next page

| Reaction name | Equation |
|---|---|
| PFK | ATP + F6P → ADP + FBP |
| PFL | COA + PYR → ACCOA + FOR |
| PGI | G6P → F6P |
| PGI_r | F6P → G6P |
| PGK | ADP + 13DPG → ATP + 3PG |
| PGK_r | ATP + 3PG → ADP + 13DPG |
| PGL | 6PGL → 6PGC |
| PGM | 3PG → 2PG |
| PGM_r | 2PG → 3PG |
| POX | PYR + Q8 → AC + CO2 + Q8H2 |
| PPC | CO2 + PEP → OAA |
| PPS | ATP + PYR → AMP + PEP |
| PTA | ACCOA → ACTP + COA |
| PTS | PEP → G6P + PYR |
| PYK | ADP + PEP → ATP + PYR |
| RPE | RU5P → XU5P |
| RPE_r | XU5P → RU5P |
| RPI | RU5P → R5P |
| RPI_r | R5P → RU5P |
| SCS | ADP + SUCCOA → ATP + COA + SUCC |
| SCS_r | ATP + COA + SUCC → ADP + SUCCOA |
| SDH | Q8 + SUCC → FUM + Q8H2 |
| SUCCt | SUCC → (Export) |
| TAL | G3P + S7P → E4P + F6P |
| TAL_r | E4P + F6P → G3P + S7P |
| THD | NADPH + NAD → NADH + NADP |
| THD_r | NADH + NADP → NADPH + NAD |
| TKT1 | R5P + XU5P → G3P + S7P |
| TKT1_r | G3P + S7P → R5P + XU5P |
| TKT2 | E4P + XU5P → F6P + G3P |
| TKT2_r | F6P + G3P → E4P + XU5P |
| TPI | G3P → DHAP |
| TPI_r | DHAP → G3P |
| | End |

Table S4: RNA expression level

| Strain | Gene | Number of copies (copies/10 ng)* | | | Mean | SD |
|---|---|---|---|---|---|---|
| | | Sample 1 | Sample 2 | Sample 3 | | |
| Vector only | gltA | N. D. | N. D. | N. D. | N/A | N/A |
| gltA | gltA | 2.11e+06 | 1.06e+06 | 1.51e+06 | 1.56e+06 | 4.30e+05 |
| Vector only | fbp | N. D. | N. D. | N. D. | N/A | N/A |
| fbp | fbp | 9.09e+05 | 8.99e+05 | 1.01e+06 | 9.41e+05 | 5.19e+04 |
| Vector only | aceA | N. D. | N. D. | N. D. | N/A | N/A |
| aceA | aceA | 3.19e+06 | 3.08e+06 | 2.41e+06 | 2.89e+06 | 3.45e+05 |
| Vector only | ldhA | 5.62e+02 | 8.26e+02 | 7.34e+02 | 7.08e+02 | 1.09e+02 |
| ldhA | ldhA | 2.12e+06 | 2.48e+06 | 9.74e+05 | 1.86e+06 | 6.41e+05 |
| Vector only | maeA | N. D. | N. D. | N. D. | N/A | N/A |
| maeA | maeA | 4.98e+05 | 4.14e+05 | 5.28e+05 | 4.80e+05 | 4.83e+04 |
| Vector only | maeB | N. D. | N. D. | N. D. | N/A | N/A |
| maeB | maeB | 2.13e+03 | 4.74e+03 | 2.59e+03 | 3.15e+03 | 1.14e+03 |
| Vector only | pckA | N. D. | N. D. | N. D. | N/A | N/A |
| pckA | pckA | 7.01e+05 | 8.53e+05 | 9.15e+05 | 8.23e+05 | 9.02e+04 |
| Vector only | ppc | N. D. | N. D. | N. D. | N/A | N/A |
| ppc | ppc | 3.52e+02 | 4.77e+02 | 2.24e+02 | 3.51e+02 | 1.03e+02 |
| Vector only | pps | N. D. | N. D. | N. D. | N/A | N/A |
| pps | pps | 1.09e+06 | 4.40e+05 | 6.11e+05 | 7.12e+05 | 2.73e+05 |
| Vector only | pta | N. D. | N. D. | N. D. | N/A | N/A |
| pta | pta | 2.91e+05 | 2.25e+05 | 1.24e+06 | 5.85e+05 | 4.63e+05 |

* Per total RNA 10 ng
N.D.: Not detected
N/A: Not applicable
SD: Standard deviation

Table S5: Equations in figures and corresponding reaction names

| Equation in figures | Reaction name |
|---|---|
| GLC → G6P | PTS |
| G6P → F6P | PGI |
| F6P → G6P | PGI_r |
| F6P → FBP | PFK |
| FBP → F6P | FBP |
| FBP → DHAP + G3P | ALD |
| DHAP + G3P → FBP | ALD_r |
| G3P → DHAP | TPI |
| DHAP → G3P | TPI_r |
| G3P → 3PG | GAPDH |
| 3PG → G3P | GAPDH_r |
| 3PG → PEP | PGM |
| PEP → 3PG | PGM_r |
| PEP → PYR | PYK |
| PYR → PEP | PPS |
| PYR → ACCOA + FOR | PFL |
| PYR → ACCOA | PDH |

Continued on next page

| Equation in figures | Reaction name |
|---|---|
| G6P → 6PGC | G6PDH |
| 6PGC → RU5P | GND |
| RU5P → XU5P | RPE |
| XU5P → RU5P | RPE_r |
| RU5P → R5P | RPI |
| R5P → RU5P | RPI_r |
| XU5P + R5P → G3P + S7P | TKT1 |
| G3P + S7P → XU5P + R5P | TKT1_r |
| S7P + G3P → E4P + F6P | TAL |
| E4P + F6P → S7P + G3P | TAL_r |
| XU5P + E4P → F6P + G3P | TKT2 |
| F6P + G3P → XU5P + E4P | TKT2_r |
| ACCOA + OAA → CIT | CS |
| CIT → ICT | ACN1 |
| ICT → CIT | ACN1_r |
| ICT → AKG | ICDH |
| AKG → SUCC | SCS |
| SUCC → FUM | SDH |
| FUM → SUCC | FRD |
| FUM → MAL | FUM |
| MAL → FUM | FUM_r |
| MAL → OAA | MDH |
| OAA → MAL | MDH_r |
| MAL → PYR | ME1 |
| ICT → GLX + SUCC | ICL |
| ACCOA + GLX → MAL | MS |
| PEP → OAA | PPC |
| OAA → PEP | PCK |
| ACCOA → AC | PTA |
| PYR → LAC | LDH |
| ACCOA → ETOH | ALDH |
| Import → GLC | GLCt |
| AC → (Export) | ACt |
| SUCC → (Export) | SUCCt |
| FOR → (Export) | FORt |
| LAC → (Export) | LACt |
| ETOH → (Export) | ETOHt |
| | End |

Table S6: Positivity confidence values calculated by BayesianSSA with the prior distribution to increase the succinate export flux.

| Perturbed reaction | Positivity confidence value |
|---|---|
| CS | 9.1270e-01 |
| FRD | 8.8220e-01 |
| FUM_r | 8.8220e-01 |
| NDH | 8.6890e-01 |
| MDH | 8.4000e-01 |
| | Continued on next page |

| Perturbed reaction | Positivity confidence value |
|---|---|
| SUCCt | 8.2650e-01 |
| GLCt | 8.1820e-01 |
| NADHDH | 8.0770e-01 |
| GAPDH | 7.9140e-01 |
| ENO | 7.9140e-01 |
| PGK | 7.9140e-01 |
| PGM | 7.9140e-01 |
| ALD | 7.9140e-01 |
| TPI_r | 7.9140e-01 |
| THD_r | 7.8990e-01 |
| PTS | 7.7900e-01 |
| PPS | 7.5590e-01 |
| PPC | 7.1430e-01 |
| FDH | 7.1430e-01 |
| ICL | 6.8610e-01 |
| PGI_r | 6.1350e-01 |
| G6PDH | 6.1350e-01 |
| SCS | 6.1230e-01 |
| PDH | 5.9550e-01 |
| PFL | 5.8390e-01 |
| GND | 5.3020e-01 |
| EDD | 4.6980e-01 |
| ADK | 4.6330e-01 |
| POX | 4.5050e-01 |
| SCS_r | 3.8770e-01 |
| PGI | 3.8650e-01 |
| PFK | 3.8650e-01 |
| CYTBO | 3.2730e-01 |
| ALDH | 3.1920e-01 |
| ICDH | 3.1390e-01 |
| ATPM | 2.9310e-01 |
| CO2t | 2.8570e-01 |
| FORt | 2.8570e-01 |
| THD | 2.1010e-01 |
| ENO_r | 2.0860e-01 |
| GAPDH_r | 2.0860e-01 |
| FBP | 2.0860e-01 |
| ALD_r | 2.0860e-01 |
| PGK_r | 2.0860e-01 |
| TPI | 2.0860e-01 |
| PGM_r | 2.0860e-01 |
| PTA | 1.9030e-01 |
| PCK | 1.7380e-01 |
| MDH_r | 1.6000e-01 |
| PYK | 1.5700e-01 |
| LDH | 1.4430e-01 |
| NDH_r | 1.3110e-01 |
| ME2 | 1.2500e-01 |
| FUM | 1.1780e-01 |
| SDH | 1.1780e-01 |

Continued on next page

| Perturbed reaction | Positivity confidence value |
|---|---|
| ME1 | 9.9900e-02 |
| | End |

Table S7: Positivity confidence values calculated by BayesianSSA fitted to the pseudo dataset to increase the succinate export flux.

| Perturbed reaction | Positivity confidence value |
|---|---|
| EDD | 9.9216e-01 |
| FRD | 9.3087e-01 |
| FUM_r | 9.3087e-01 |
| NDH | 9.0215e-01 |
| SUCCt | 8.8060e-01 |
| PPC | 8.6867e-01 |
| FDH | 8.6867e-01 |
| CS | 7.6992e-01 |
| NADHDH | 7.4237e-01 |
| PFL | 7.0145e-01 |
| GAPDH | 6.9603e-01 |
| ALD | 6.9603e-01 |
| PGK | 6.9603e-01 |
| PGM | 6.9603e-01 |
| TPI_r | 6.9603e-01 |
| ENO | 6.9603e-01 |
| PGI_r | 6.4605e-01 |
| G6PDH | 6.4605e-01 |
| SCS | 6.1495e-01 |
| THD_r | 6.0471e-01 |
| ICL | 5.8966e-01 |
| PYK | 5.6871e-01 |
| MDH | 5.4890e-01 |
| POX | 5.4673e-01 |
| PPS | 5.4563e-01 |
| ALDH | 5.2737e-01 |
| PDH | 5.2213e-01 |
| ADK | 4.8118e-01 |
| MDH_r | 4.5110e-01 |
| ICDH | 4.1034e-01 |
| ATPM | 3.9895e-01 |
| THD | 3.9529e-01 |
| SCS_r | 3.8505e-01 |
| PGI | 3.5395e-01 |
| PFK | 3.5395e-01 |
| ME1 | 3.4051e-01 |
| ME2 | 3.4038e-01 |
| LDH | 3.1621e-01 |
| TPI | 3.0397e-01 |
| FBP | 3.0397e-01 |
| PGK_r | 3.0397e-01 |
| | Continued on next page |

| Perturbed reaction | Positivity confidence value |
|---|---|
| ALD_r | 3.0397e-01 |
| GAPDH_r | 3.0397e-01 |
| ENO_r | 3.0397e-01 |
| PGM_r | 3.0397e-01 |
| CYTBO | 2.7508e-01 |
| GLCt | 2.2727e-01 |
| PCK | 1.7239e-01 |
| CO2t | 1.3133e-01 |
| FORt | 1.3133e-01 |
| PTA | 1.2914e-01 |
| NDH_r | 9.7846e-02 |
| FUM | 6.9131e-02 |
| SDH | 6.9131e-02 |
| PTS | 3.2294e-02 |
| GND | 7.8447e-03 |
| | End |

Table S8: Positivity confidence values calculated by BayesianSSA fitted to the real dataset to increase the succinate export flux.

| Perturbed reaction | Positivity confidence value |
|---|---|
| GLCt | 9.8965e-01 |
| ALD | 9.8709e-01 |
| TPI_r | 9.8709e-01 |
| GAPDH | 9.8709e-01 |
| PGM | 9.8709e-01 |
| ENO | 9.8709e-01 |
| PGK | 9.8709e-01 |
| PTS | 9.8381e-01 |
| MDH | 9.7380e-01 |
| CS | 9.6357e-01 |
| THD_r | 9.4574e-01 |
| ICL | 9.4262e-01 |
| NDH | 9.1748e-01 |
| CO2t | 9.0747e-01 |
| FORt | 9.0747e-01 |
| FRD | 9.0544e-01 |
| FUM_r | 9.0544e-01 |
| PPS | 8.3850e-01 |
| NADHDH | 8.0402e-01 |
| SUCCt | 7.9912e-01 |
| PCK | 7.9559e-01 |
| GND | 5.7501e-01 |
| PFL | 5.6591e-01 |
| SCS | 5.6194e-01 |
| G6PDH | 5.3732e-01 |
| PGI_r | 5.3732e-01 |
| ADK | 5.0079e-01 |
| | Continued on next page |

| Perturbed reaction | Positivity confidence value |
|---|---|
| PDH | 4.6290e-01 |
| PFK | 4.6268e-01 |
| PGI | 4.6268e-01 |
| SCS_r | 4.3806e-01 |
| EDD | 4.2499e-01 |
| POX | 2.6887e-01 |
| ATPM | 1.6815e-01 |
| CYTBO | 1.6631e-01 |
| PTA | 1.2527e-01 |
| FUM | 9.4564e-02 |
| SDH | 9.4564e-02 |
| PPC | 9.2534e-02 |
| FDH | 9.2534e-02 |
| NDH_r | 8.2516e-02 |
| ALDH | 7.7361e-02 |
| ICDH | 5.7378e-02 |
| THD | 5.4258e-02 |
| MDH_r | 2.6197e-02 |
| PGK_r | 1.2909e-02 |
| TPI | 1.2909e-02 |
| PGM_r | 1.2909e-02 |
| ALD_r | 1.2909e-02 |
| FBP | 1.2909e-02 |
| GAPDH_r | 1.2909e-02 |
| ENO_r | 1.2909e-02 |
| ME2 | 1.1993e-02 |
| PYK | 9.8488e-03 |
| LDH | 9.6581e-03 |
| ME1 | 1.1440e-03 |
| | End |

Table S9: Positivity confidence values calculated by BayesianSSA fitted to the real dataset using log normal distributions with random parameters to increase the succinate export flux.

| Perturbed reaction | Positivity confidence value |
|---|---|
| GLCt | 9.6239e-01 |
| PTS | 9.5010e-01 |
| ICL | 9.3958e-01 |
| GAPDH | 9.3824e-01 |
| PGK | 9.3795e-01 |
| PGM | 9.3608e-01 |
| ALD | 9.3424e-01 |
| ENO | 9.3101e-01 |
| TPI_r | 9.3037e-01 |
| MDH | 9.0591e-01 |
| CO2t | 8.9606e-01 |
| FORt | 8.9499e-01 |
| CS | 8.8738e-01 |

Continued on next page

| Perturbed reaction | Positivity confidence value |
|---|---|
| FRD | 8.8063e-01 |
| FUM_r | 8.7741e-01 |
| THD_r | 8.6143e-01 |
| PCK | 8.4588e-01 |
| NDH | 8.3607e-01 |
| SUCCt | 8.3580e-01 |
| PPS | 7.4961e-01 |
| NADHDH | 6.3783e-01 |
| SCS | 6.1182e-01 |
| GND | 6.0631e-01 |
| PFL | 5.9157e-01 |
| PGI | 5.4369e-01 |
| PFK | 5.3852e-01 |
| ADK | 5.2466e-01 |
| PDH | 4.7393e-01 |
| G6PDH | 4.4985e-01 |
| PGI_r | 4.4899e-01 |
| EDD | 3.9199e-01 |
| SCS_r | 3.8119e-01 |
| ATPM | 3.2834e-01 |
| POX | 2.9159e-01 |
| CYTBO | 2.3841e-01 |
| NDH_r | 1.6154e-01 |
| ALDH | 1.5794e-01 |
| PTA | 1.3126e-01 |
| THD | 1.2863e-01 |
| FUM | 1.1726e-01 |
| SDH | 1.1706e-01 |
| FDH | 9.8546e-02 |
| PPC | 9.5287e-02 |
| MDH_r | 8.5864e-02 |
| ICDH | 5.9483e-02 |
| ENO_r | 5.1211e-02 |
| PGK_r | 4.9560e-02 |
| GAPDH_r | 4.6104e-02 |
| PGM_r | 4.4578e-02 |
| PYK | 4.4059e-02 |
| TPI | 4.2607e-02 |
| LDH | 4.1577e-02 |
| ALD_r | 3.8872e-02 |
| FBP | 3.5333e-02 |
| ME2 | 3.4725e-02 |
| ME1 | 9.7625e-03 |
| | End |
